# Dynamic Nanoparticle Assembly-Based Biomedical Microrobots

**DOI:** 10.64898/2026.07.30.741696

**Authors:** Lukas Hertle, Hao Ye, Hyeon Ko, Carlos Franco, Valentin Gantenbein, Derick Sivakumaran, Ishika Paul, Minsoo Kim, Andrea Veciana, Laura Baraldi, Zhengwei Tan, Fabian C. Landers, Pascal Theiler, Pere Bruna, Minghan Hu, Yongfeng Mei, Eneko Garaio, Alberto López-Ortega, Josep Puigmarti-Luis, Miriam Weisskopf, Xiang-Zhong Chen, Bradley J. Nelson, Salvador Pané

## Abstract

Precise drug delivery within anatomically complex tissues demands systems capable of both active navigation and deep tissue access, properties that have remained difficult to reconcile in existing nanocarriers and microrobots. Here we introduce Dynabots, a dynamic microrobotic assembly constructed from multifunctional nanoparticles covalently linked by thermally cleavable molecular connectors. This nanoparticle-rich architecture enables the integration of magnetic, imaging, and therapeutic components while preserving a high content of functional material. Collective assembly imparts enhanced magnetic responsiveness and maneuverability, enabling controlled navigation through tortuous biological environments. Upon exposure to mild thermal stimuli, the assemblies undergo programmed disassembly, releasing individual nanoparticles that can diffuse through tissue for localized therapeutic action. We establish the programmable transitions, biocompatibility, and therapeutic efficacy of this process across *in vitro* and *in vivo* models, including real-time fluoroscopic guidance within anatomically realistic phantoms and live rodent and porcine systems. By integrating magnetic control, reconfigurable architecture, and stimulus-triggered disassembly, Dynabots unite navigational precision with tissue permeability, providing a versatile platform for adaptive and deep-tissue drug delivery.

## Introduction

Targeted delivery is key for treating diseases located in anatomically hard-to-reach areas, where conventional systemic administration fails to achieve therapeutic efficacy without inducing significant off-target toxicity.(*1*, *2*) Examples include brainstem gliomas, intraventricular tumors, and deep-seated pancreatic malignancies, where physiological barriers such as the blood-brain barrier and complex vascular architecture severely limit drug access. While nanomedicine has made significant strides in developing biocompatible drug carriers, most nanoparticle-based delivery systems rely on passive accumulation or ligand-mediated targeting and lack active steering capabilities.(*3–5*) As a result, drug accumulation at the intended site remains inefficient and inconsistent - leading to the striking reality that over 99% of systemically administered therapeutics are wasted.(*6*)

Magnetic microrobots have emerged as a promising solution for targeted delivery by enabling active navigation through externally applied magnetic fields, allowing precise access to deep or complex anatomical regions that are otherwise unreachable via passive means.(*7–10*) However, a fundamental challenge remains unresolved: the mismatch between the physical requirements for magnetic navigation and those for effective tissue penetration and therapeutic action. Larger microrobots can be steered over long distances but struggle to infiltrate dense or heterogeneous tissues such as solid tumors. Conversely, magnetic nanoparticles small enough to diffuse through tissue microenvironments cannot be effectively maneuvered in vivo due to insufficient magnetic responsiveness. Incorporating magnetic nanoparticles into polymer matrices can improve their collective magnetic response. However, the achievable loading may involve trade-offs with the mechanical properties, structural integrity, and processability of the composite.

To bridge the gap between the magnetic controllability of microrobots and the tissue-penetrating ability of nanoparticles, we introduce Dynabots, a dynamic, modular microrobotic platform. The term “dynamic” reflects their structural transitions between assembled and disassembled states.(*11*, *12*) In their assembled form, Dynabots exhibit enhanced magnetic responsiveness and fluidic stability, enabling efficient locomotion and robust control during navigation, while simultaneously encapsulating drug molecules to prevent premature release. Upon reaching the target lesion, the use of mild thermal stimuli can be applied to trigger on-demand disassembly through thermally cleavable linkers, releasing individual nanoparticles that diffuse deeply into tissue and achieve localized therapeutic delivery.

Dynabots are built on a modular assembly design, allowing for the incorporation of diverse functional nanoparticles, including magnetic components for actuation, imaging agents for fluoroscopic tracking, and therapeutic payloads for treatment (Fig. 1a). Unlike conventional polymer composites, this architecture requires only a small amount of nonfunctional material to hold the nanoparticles together, thereby preserving a high proportion of active components. The building blocks are covalently linked through surface-modified chains containing polyethylene glycol (PEG), which provides excellent biocompatibility and helps minimize immunogenicity. Crucially, the PEG linkers are functionalized with maleimide-furan Diels-Alder adducts, enabling thermally induced retro-Diels-Alder cleavage.(*13*) This chemical feature ensures precise and controllable disassembly of Dynabots under clinically relevant magnetothermal or photothermal conditions (Fig. 1b,c). Beyond their dynamic responsiveness, Dynabots exhibit high compatibility with existing microrobotic manipulation strategies.(*14–18*) They can be configured into small-scale clusters, suitable for swarm-based magnetic actuation, leveraging cooperative behavior for enhanced navigation and surface adaptability. Alternatively, Dynabots can form monolithic, magnetically steerable microrobots, capable of long-range propulsion and real-time imaging in complex anatomical settings (Fig. 1d,e and fig. S1). This dual-mode compatibility offers a functional enhancement over conventional swarm or monolithic systems. This flexibility allows Dynabots to bridge the gap between microrobotics and nanomedicine, enabling precision therapy through structural reconfiguration, active navigation, and imaging-guided delivery - all within a single integrated platform.

**Fig. 1.**
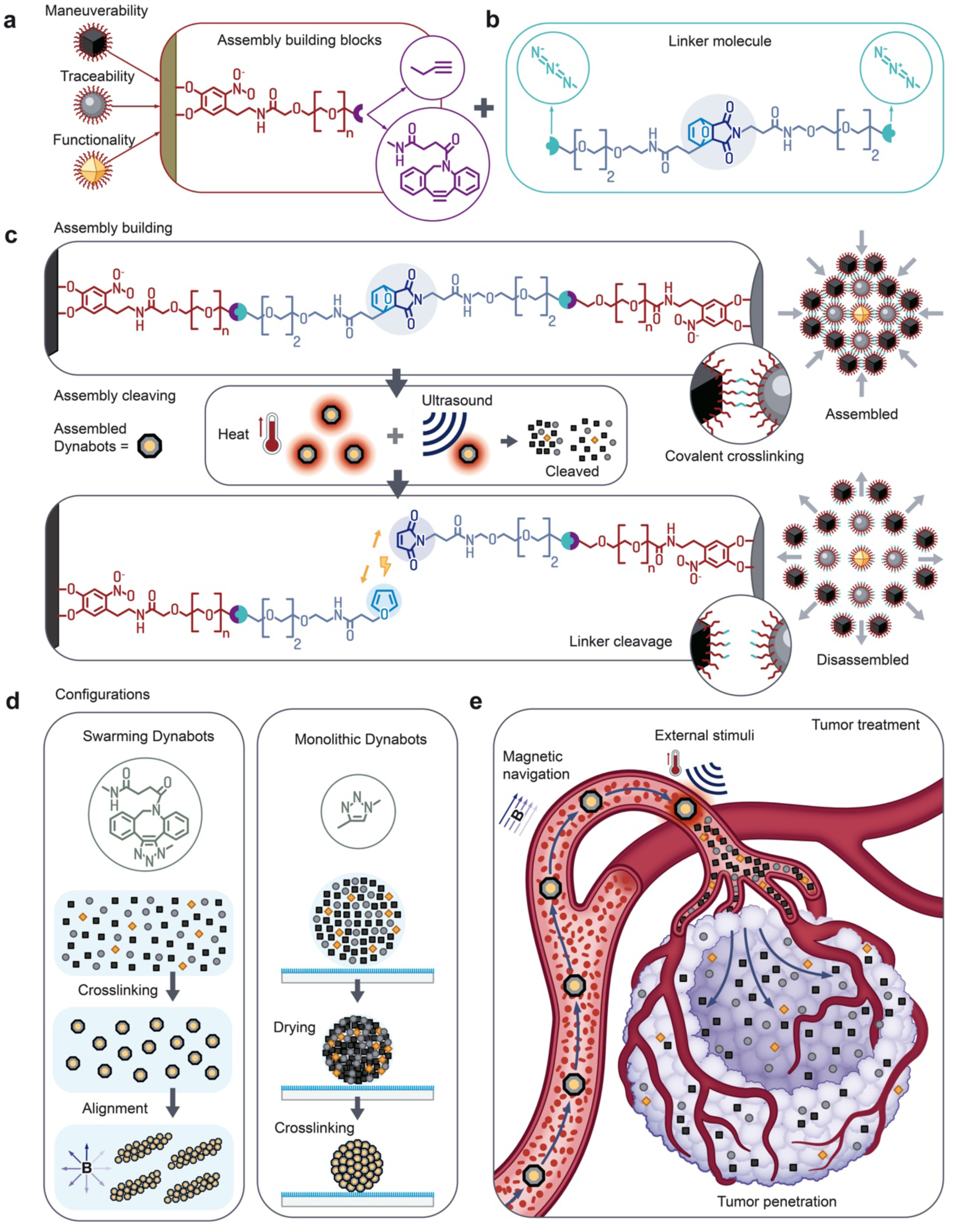
Conceptual illustration. **a**, Schematic representation of the nanoparticle building blocks and their functional surface ligands. **b,** Chemical structure of the synthesized thermo-responsive crosslinker. **c,** Conceptual illustration of nanoparticle assembly formation via covalent crosslinking and its disassembly triggered by heating. **d,** Schematic showing programmable microrobot swarm formation and monolithic structure fabrication. **e,** Schematic illustration of different microrobotic delivery strategies for targeted therapy.

### Functional nano building blocks and their dynamic assembly

The fabrication of Dynabots began with the synthesis of multiple functional nanoparticle components, each designed to impart essential microrobotic capabilities (Fig. 1a). Zinc-substituted iron oxide (Zn_x_Fe_3-x_O₄) nanoparticles were employed to provide magnetic maneuverability, leveraging the Zn(II)-induced suppression of antiferromagnetic superexchange interactions to enhance magnetic responsiveness of the nanoparticles.(*19–21*) Tantalum nanoparticles were incorporated to enable fluoroscopic traceability under X-ray,(*22*) while UiO-66-NH_2_ metal-organic frameworks were selected as drug carriers owing to their high drug-loading capacity.(*23*) All particles were surface-modified with short polyethylene glycol (PEG) chains bearing either terminal alkyne or cyclooctyne groups, following established protocols to facilitate covalent crosslinking while also ensuring high biocompatibility and colloidal stability in physiological media.(*24–26*) High-angle annular dark-field scanning transmission electron microscopy (HAADF-STEM) confirmed the successful synthesis of highly crystalline monodisperse cubic Zn_x_Fe_3-x_O₄ nanoparticles with an average edge length of 17.9 ± 1.7 nm and a width of 25.6 ± 1.9 nm (Fig. 2a,c). Energy-dispersive X-ray (EDX) mapping revealed a homogeneous distribution of Fe, Zn, and O elements within individual particles (Fig. 2a). X-ray diffraction (XRD) and Mössbauer spectroscopy (fig. S2) confirmed the formation of an inverse spinel phase, with a crystalline domain size of ∼ 15 nm as calculated from the (311) peak using the Scherrer equation. Surface modification was conducted in two steps: (i) replacement of native oleic acid ligands with nitrodopamine, a chelator with high Fe^2+^ affinity; and (ii) conjugation of PEG chains terminated with either alkyne or dibenzocyclooctyne (DBCO) groups via N-hydroxysuccinimide (NHS) chemistry targeting surface amines (Fig. 2b). The successful synthesis of nitrodopamine and subsequent surface functionalization were confirmed using proton nuclear magnetic resonance (¹H-NMR), Fourier-transform infrared spectroscopy (FTIR), thermogravimetric analysis (TGA), and dynamic light scattering (DLS) (figs. S3 and S4). Ligand grafting densities, calculated from TGA mass loss, indicated an average of ∼ 6.0 nitrodopamine anchors, ∼ 0.6 Alk-PEG chains, and ∼ 0.3 DBCO-PEG chains per nm^2^ surface area, respectively, ensuring sufficient surface reactivity for subsequent assembly. The same functionalization strategy was applied to UiO-66-NH_2_ particles and commercial tantalum nanoparticles (see Methods) and equally confirmed by FTIR and TGA analysis (figs. S5 and S6). Vibrating sample magnetometry (VSM) demonstrated that Zn_x_Fe_3-x_O_4_ nanoparticles exhibited high saturation magnetization (∼90 emu g_IO_⁻¹) and strong magnetic responsiveness, achieving ∼73% of saturation at 30 mT (∼66 emu g_IO_⁻¹) (Fig. 2d). Notably, alternating current hysteresis measurements showed no significant change in specific absorption rate (SAR) before and after PEG modification, indicating preserved heating efficiency and a Néel-dominated relaxation (Fig. 2e and fig. S7).

**Fig. 2.**
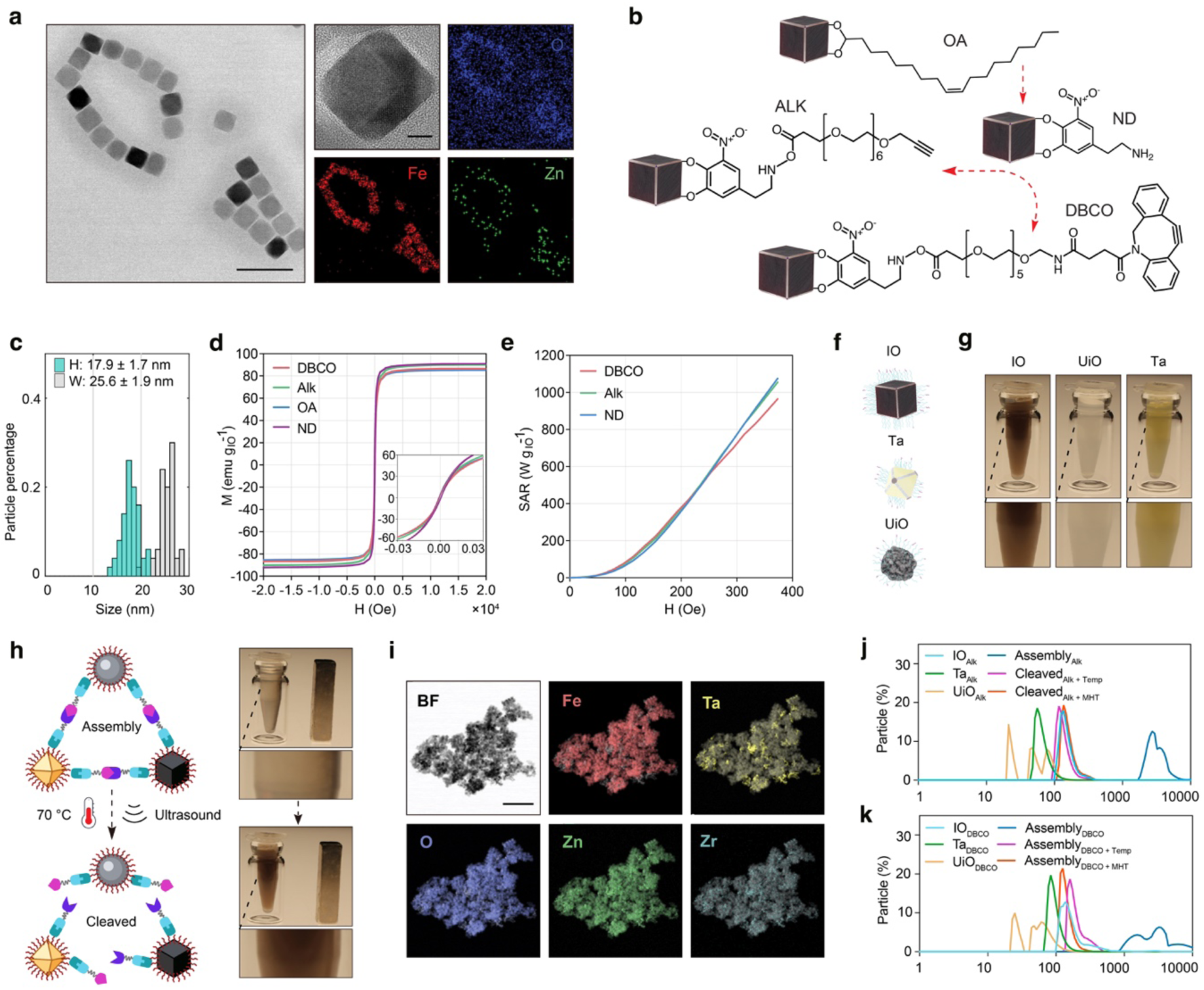
Material characterization and Dynabot assembly/disassembly. **a**, TEM images, a high-magnification image and EDX elemental maps of zinc-ferrite nanoparticles. Scale bars, 50 nm (left) and 5 nm (middle). **b,** Schematic illustration of the surface modification protocol. **c,** Size distribution of zinc-ferrite nanoparticles (n = 50). **d,** Magnetization curves of zinc-ferrite nanoparticles functionalized with different ligand molecules. **e,** Field-dependent specific absorption rate (SAR) values of ligand-modified zinc-ferrite nanoparticles at 480 kHz. **f,** Schematic illustration of the three nanoparticle building blocks and **g,** digital images of the dispersion states of the nanoparticles in deionized water. **h,** Illustration of the assembly process of the three nanoparticle types and the triggered disassembly, with corresponding digital images of the assembled clusters and cleaved structures in deionized water. **i,** Bright-field SEM image and EDX elemental maps of the assembled Dynabots. Scale bar, 200 nm. **j,** Hydrodynamic diameters of the functionalized nanoparticles, the assemblies and the disassembled products for the alkyne-terminated and **k,** DBCO-terminated systems (n = 3).

The thermo-responsive bis-azide crosslinker was synthesized based on a maleimide-furan Diels-Alder (DA) adduct bearing two triethylene glycol chains. This molecule undergoes retro-Diels-Alder cleavage at temperatures above 60 °C,(*13*) allowing reversible linkage between nanoparticles. The PEG backbone imparts water solubility, while azide end groups facilitate copper-catalyzed or copper-free click reactions with alkyne- or cyclooctyne-functionalized nanoparticles. The chemical identity and cleavability of the crosslinker were confirmed by ^1^H-NMR and FTIR (fig. S8), and thermal cleavage was further verified by ^1^H-NMR after heating at 70 °C, which showed near-complete retro-DA reaction (fig. S9).

Dynamic nanoparticle assemblies were obtained by mixing the functionalized particles with the synthesized bis-azide crosslinkers, which allowed them to react via either copper-catalyzed or copper-free click reactions, depending on the terminal groups (Fig. 2f-h and Movie S1). Scanning electron microscopy (SEM) and EDX mapping confirmed the formation of heterogeneous micrometer-sized clusters incorporating all three particle types (Fig. 2i). The average assembly size was tuneable by varying the particle-to-crosslinker ratio, as reflected in DLS measurements (fig. S10). Small-angle X-ray scattering (SAXS) analysis revealed radii of gyration of ∼42.2 Å (DBCO-terminated) and ∼43.6 Å (alkyne-terminated) and estimated distances between two scattering points of ∼ 126.4 (DBCO-terminated) and 120.2 Å (alkyne-terminated), consistent with disordered internal structure and interparticle distances close to the expected linker length (∼15 nm, fig. S11).(*27–30*) Assuming this interparticle spacing, we numerically simulated the SAR and AC hysteresis loops of single IO particles and assembled Dynabots to determine whether assembly alters their dynamic magnetic properties. The simulated SAR and AC hysteresis loops of assembled Dynabots were almost identical to those of dispersed nanoparticles, indicating similar heating mechanisms and efficiencies in both environments (fig. S12). Exposure of the assembled clusters to a high-frequency alternating magnetic field (AMF; 20 mT, 510 kHz) led to disassembly (Fig. 2h). DLS measurements showed that the hydrodynamic diameters of the disassembled products were slightly larger (∼100 nm) than those of pristine individual nanoparticles and were consistent between products cleaved by MHT (Cleaved MHT) and those cleaved by bulk heating to 60 °C (Cleaved Temp). This cleavage behavior was consistent for both DBCO- and alkyne-linked assemblies (Fig. 2j,k). This slight size increase likely reflects partial nanoparticle aggregation (e.g., dimers or trimers) influenced by the linker end groups. Nevertheless, the disassembled nanoparticles remained colloidally stable in aqueous solution, as confirmed by visual dispersion and stability data. Further stability studies in phosphate buffered saline (PBS) showed that the particles maintained a hydrodynamic diameter of ∼350 nm over time (fig. S13), highlighting their suitability for *in vivo* use. Importantly, pulsed AMF stimulation was able to trigger disassembly at a bulk solution temperature of just 42 °C, which is below cytotoxic thresholds, suggesting that localized particle surface heating was sufficient to induce linker cleavage without significantly increasing surrounding temperature.(*31–33*)

### *In vitro* biocompatibility and therapeutic efficacy

To evaluate biocompatibility, NIH/3T3 embryonic fibroblast cells were incubated for 72 hours with Dynabot components at various stages: individual nanoparticles prior to assembly, intact assemblies, and post-disassembly products. Live/dead staining and MTT assays revealed high biocompatibility of all tested samples at a concentration of 150 mg L⁻¹, with over 80% cell viability observed in each condition (Fig. 3a-c and fig. S14).(*34*) No statistically significant difference in viability was detected between assemblies formed via alkyne- or DBCO-modified nanoparticles, whether intact or disassembled. While a slight decline in viability was noted at higher concentrations, all groups maintained over 80% viability even at 150 mg L⁻¹ (Fig. 3c and fig. S15), indicating excellent cytocompatibility of the Dynabot platform in vitro.

**Fig. 3.**
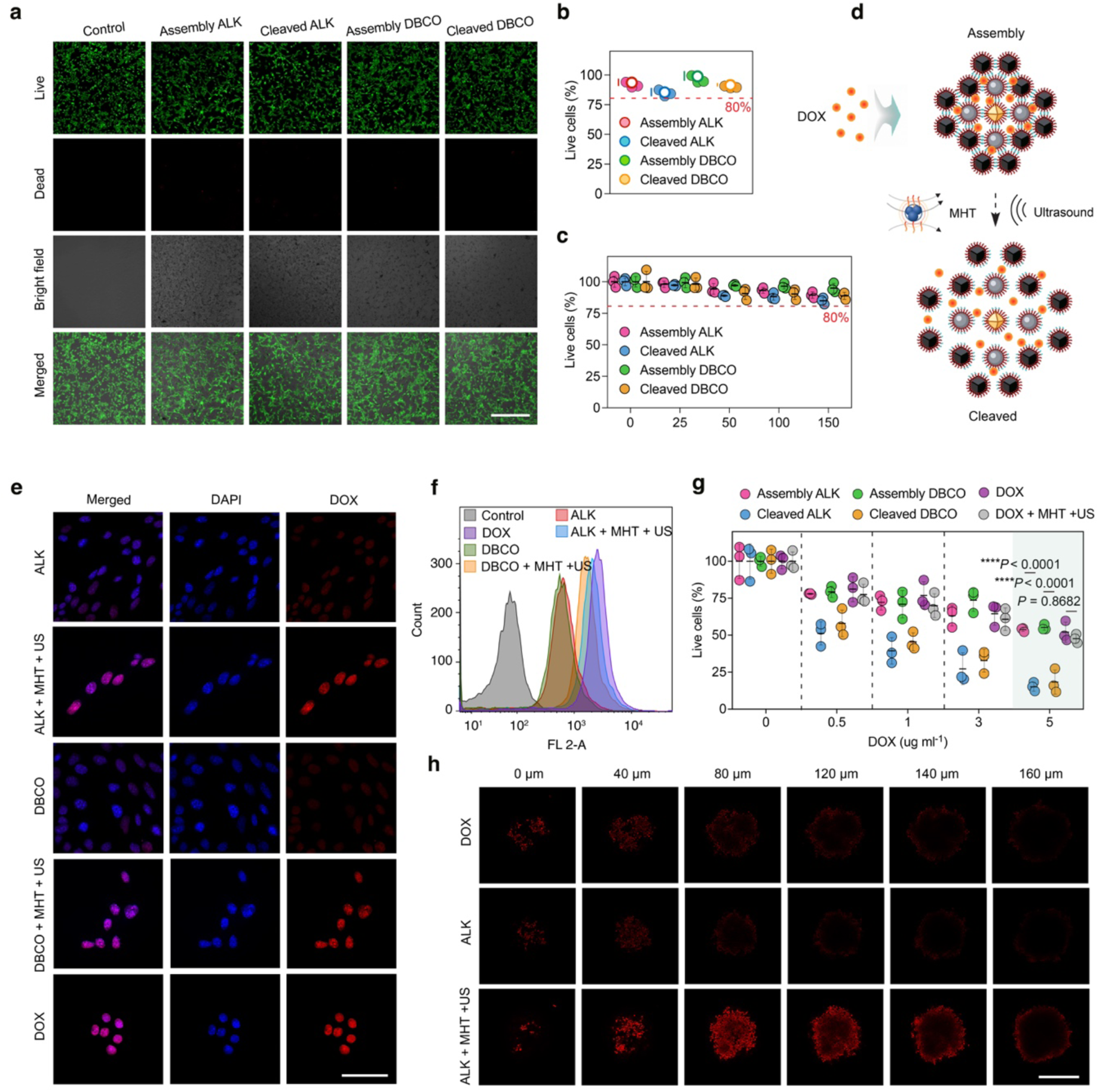
In vitro biocompatibility, triggered drug release and tumor penetration of Dynabots. **a**, Live/dead staining of NIH/3T3 cells exposed for 72 h to alkyne-based assemblies, cleaved alkyne-based assemblies, DBCO-based assemblies and cleaved DBCO-based assemblies. Scale bar, 180 μm. **b,** Quantification of live cells from the live/dead assay in a (n = 3). **c,** MTT assays of B16-F10 cells exposed to alkyne-based assemblies, cleaved alkyne-based assemblies, DBCO-based assemblies and cleaved DBCO-based assemblies at the indicated concentrations (n = 3). **d,** Schematic illustration of DOX loading into Dynabots and triggered release upon magnetic hyperthermia (MHT) and ultrasound-induced disassembly. **e,** Confocal images of B16-F10 cells incubated with the indicated samples for 2 h with and without MHT and ultrasound stimulation. Scale bar, 40 μm. **f,** Flow cytometry of intracellular DOX in B16-F10 cells treated with the indicated samples. **g,** Cytotoxicity of B16-F10 cells exposed for 24 h to free DOX, DOX-loaded alkyne-based assemblies, cleaved alkyne-based assemblies, DOX-loaded DBCO-based assemblies and cleaved DBCO-based assemblies at the indicated DOX concentrations (n = 3). **h,** Confocal z-stack images of 3D B16-F10 tumour spheroids after incubation with free DOX or DOX-loaded alkyne-based assemblies for 4 h with and without MHT and ultrasound stimulation. Scale bar, 200 μm.

To investigate the therapeutic potential, we next loaded the assembled Dynabots with doxorubicin (DOX) through absorption which was confirmed by the recorded release profile at a pH 5.5 (fig. S16), before monitoring the cellular uptake of the drug molecules. When incubated with intact Dynabots, B16-F10 melanoma cells exhibited limited DOX internalization. However, upon exposure to magnetic hyperthermia (MHT) and ultrasound, which triggered the disassembly of Dynabots, significantly enhanced intracellular DOX accumulation was observed by confocal fluorescence imaging (Fig. 3d,e). Flow cytometry further validated these results (Fig. 3f), supporting the hypothesis that structural disassembly reduces particle size, thereby improving tumor cell penetration and drug delivery efficiency.(*35–37*)

This triggered release strategy enhances therapeutic efficacy, as evidenced by the cell viability assays (Fig. 3g). At a low DOX concentration of 5 μg mL⁻¹, over 80% of cancer cells were eliminated when treatment was combined with MHT-induced disassembly. In contrast, minimal cytotoxicity was observed when DOX-loaded Dynabots were administered without external activation. This highlights the importance of stimulus-responsive release in enabling effective treatment. Notably, the enhanced therapeutic outcome may also result from a synergistic combination of efficient DOX release and localized hyperthermia-enhanced chemotherapeutic efficacy induced by the applied MHT.(*33*, *38*)

To evaluate performance in tissue-like environments, we employed 3D tumour spheroid models to assess DOX penetration. As shown in Fig. 3h, disassembled Dynabots enabled markedly improved drug infiltration and retention within the spheroids. Quantitative analysis revealed ∼80% higher fluorescence intensity compared to free DOX, while intact Dynabots (without disassembly) showed ∼38% lower signal intensity (fig. S17). These results demonstrate both the robust cargo-retention capability of the intact assemblies and their capacity for on-demand release, thereby minimizing premature diffusion and maximizing site-specific therapeutic delivery.

### *In vivo* therapeutic efficacy and biosafety of Dynabots

To evaluate the *in vivo* anti-tumour efficacy of Dynabots, we conducted therapeutic studies in a murine melanoma model. Given that prior *in vitro* experiments showed no significant difference in biocompatibility or drug delivery performance between alkyne- and DBCO-linked assemblies, the *in vivo* studies were carried out exclusively using alkyne-based Dynabots, in alignment with the 3R principle (Replacement, Reduction, and Refinement). As illustrated in Fig. 4a, DOX-loaded Dynabots were administered via peritumoral injection, followed by localized thermal and ultrasound stimulation of the tumour site over the first 6 days post-injection. Mice were subsequently monitored and sacrificed on day 20 for endpoint analysis. Due to the unavailability of magnetic hyperthermia equipment suitable for small animals, thermal stimulation was performed using near-infrared (NIR) light to induce photothermal heating of the magnetic nanoparticles. As shown in Fig. 4b,c, infrared thermography recorded an apparent tissue-surface temperature of ∼55 °C within 5 min under 808 nm irradiation. Because this surface measurement may underestimate the temperature at the Dynabot-rich region, we performed COMSOL-based thermal modelling of a dispersed, disc-shaped heat-generating domain beneath the skin. The simulation suggested that the local temperature within the Dynabot-rich region could reach ∼72 °C while the apparent tissue-surface temperature remained ∼55–60 °C. Because blood perfusion was neglected, this value should be interpreted as an upper-bound estimate. Nevertheless, the result supports the feasibility of local ThermoXlinker activation in vivo (fig. S18).

**Fig. 4.**
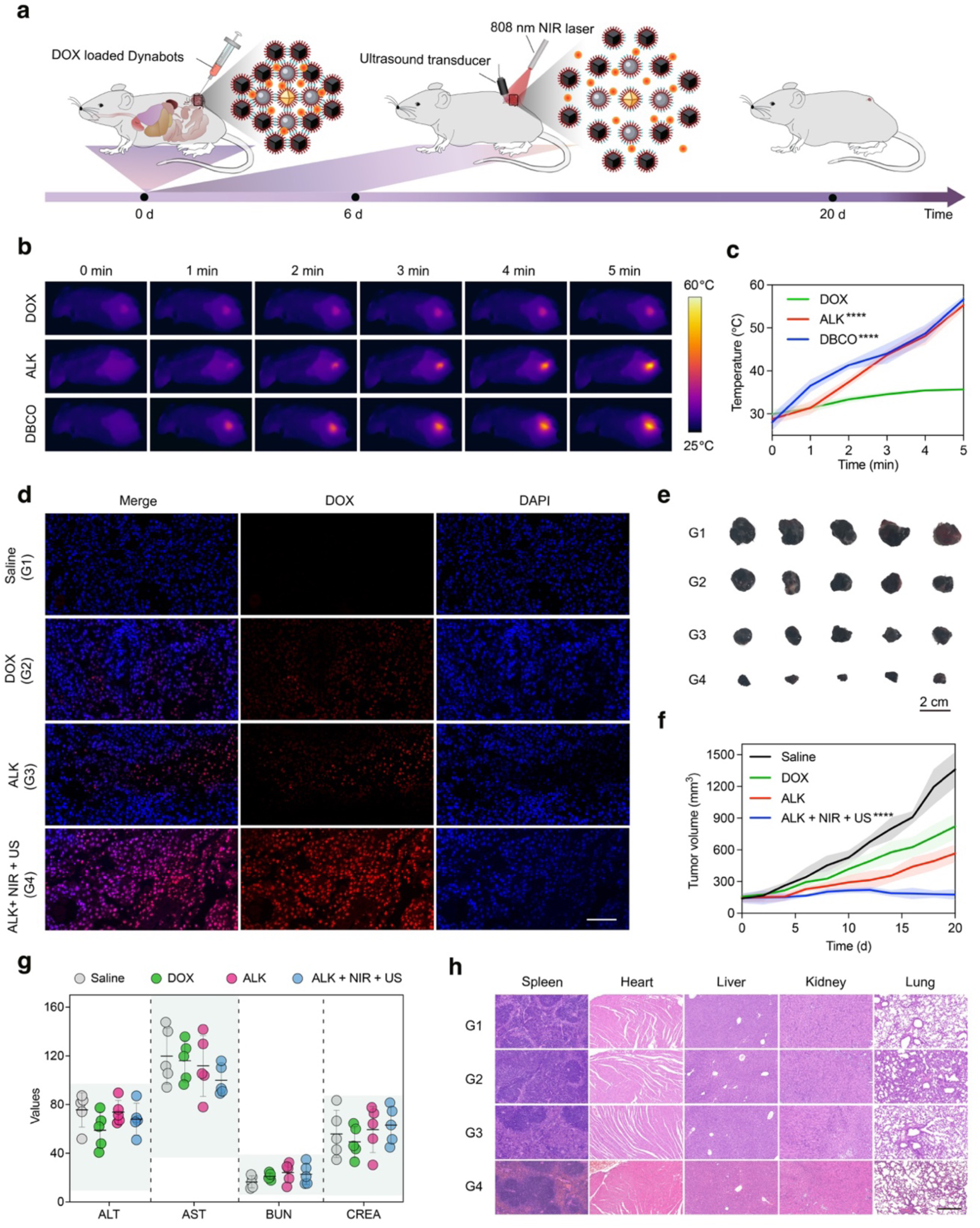
*In vivo* therapeutic efficacy and biosafety of Dynabots. **a**, Schematic illustration of the in vivo treatment protocol. DOX-loaded Dynabots were administered by peritumoral injection, followed by localized NIR and ultrasound stimulation, and animals were monitored until day 20. **b,** Representative thermal images and **c,** quantified apparent tissue-surface heating profiles of free DOX, alkyne-linked Dynabots and DBCO-linked Dynabots under 808 nm NIR irradiation (n = 5). **d,** Confocal images of tumour sections showing DOX distribution after treatment with saline (G1), free DOX (G2), DOX-loaded alkyne-linked Dynabots (G3) and DOX-loaded alkyne-linked Dynabots with NIR + US activation (G4). Scale bar, 200 μm. **e,** Representative excised tumours and **f,** tumour growth curves for the indicated treatment groups over 20 d (n = 5). Scale bar, 2 cm. **g,** Hepatorenal function parameters of blood obtained at day 20, including serum ALT, AST, BUN and CREA levels, for the indicated treatment groups (n = 5). **h,** H&E staining of major organs collected from the indicated treatment groups at day 20. Scale bar, 2 mm.

Fluorescence imaging of excised tumours revealed that Dynabots, when combined with thermal and ultrasound activation, enabled significantly enhanced DOX accumulation within tumour tissues (Fig. 4d), consistent with in vitro uptake trends. Tumour growth measurements over a 20-day period demonstrated that mice treated with activated DOX-loaded Dynabots exhibited substantial tumor regression, with an average volume reduction of ∼87% relative to untreated controls (Fig. 4e,f). This effect was markedly greater than that achieved by free DOX (∼40% reduction) or by DOX-loaded Dynabots without disassembly (∼58% reduction), highlighting the critical role of triggered structural disassembly in therapeutic enhancement.

To assess the systemic safety of Dynabot-based treatment, we measured key serum biomarkers of hepatic and renal function, including creatinine (CREA), blood urea nitrogen (BUN), alanine aminotransferase (ALT), and aspartate aminotransferase (AST). All values remained within physiological reference ranges (Fig. 4g), indicating no signs of hepatic or renal toxicity. Furthermore, histological analysis of major organs (heart, liver, spleen, lung, and kidney) revealed no significant morphological abnormalities across all treatment groups (Fig. 4h). Consistent with these findings, body weights remained stable throughout the study (fig. S19), further supporting the systemic biosafety of the platform.

Collectively, these results demonstrate that Dynabots offer potent and site-specific tumour suppression when combined with external triggering, while exhibiting minimal off-target toxicity, underscoring their potential for safe and effective clinical translation.

### Maneuverability and navigation of Dynabot swarms

To evaluate the navigation capabilities of the micrometre-sized Dynabots as programmable delivery swarms, we investigated their collective maneuverability in complex physiological environments. Swarm-based delivery has recently attracted growing interest due to its intrinsic adaptability to environmental fluctuations, improved functional redundancy, and enhanced resilience against perturbations.(*14–16*, *39–41*) Such swarm behaviour can be modulated through external control over magnetic field parameters (Fig. 5a).

**Fig. 5.**
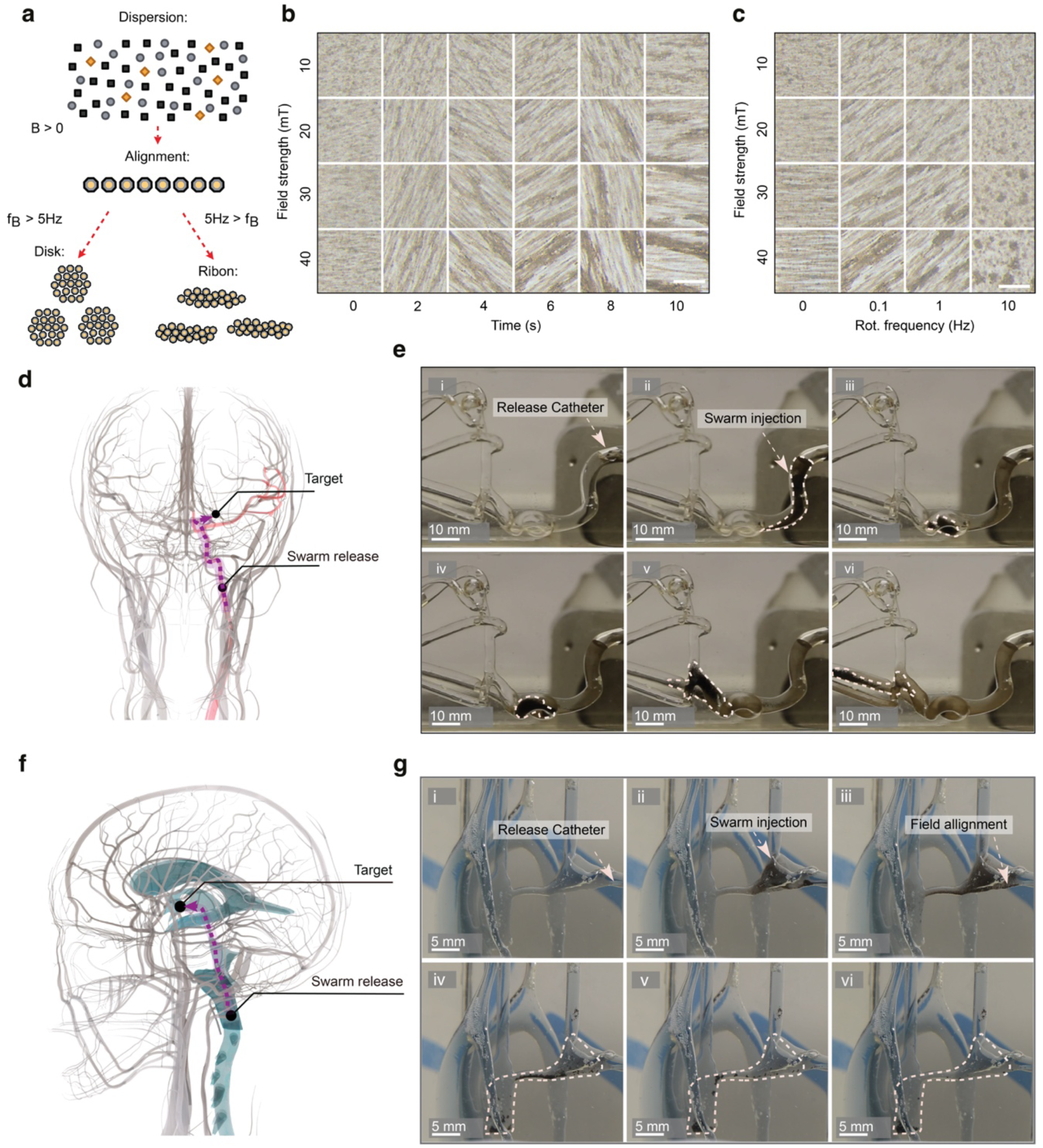
Dynabot swarm formation and navigation within complex biological architectures. **a**, Schematic illustration of the field-dependent morphological transitions of Dynabots from dispersed particles to aligned structures and then to disk-like or ribbon-like swarms under different magnetic actuation conditions. **b,** Time-lapse optical images showing swarm formation at different magnetic field strengths. Scale bars, 400 μm. **c,** Optical images showing the morphology of the swarms as a function of rotating-field frequency and magnetic field strength. Scale bars, 400 μm. **d,** Schematic overview of human cerebral vasculature with highlighted region corresponding to the middle cerebral artery (MCA) phantom model. **e,** Time-lapse images of Dynabot injection and magnetically guided propulsion through the MCA phantom. **f,** Schematic illustration of the human ventricular system with highlighted regions corresponding to the phantom model. **g,** Time-lapse images showing Dynabot navigation through a human brain ventricular phantom under rotating magnetic field actuation.

A systematic study of swarm formation revealed that increasing magnetic field strength promoted stronger dipolar interactions, leading to tighter inter-assembly alignment and more cohesive collective motion (Fig. 5b and fig. S20). Moreover, rotational frequency of the magnetic field emerged as a key determinant of swarm morphology. At low frequencies, rotating ribbon-like formations were observed, while higher frequencies induced spherical, compact swarms (Fig. 5c). The applied field’s magnitude and frequency govern the magnetic dipole interactions and alignment forces, thereby regulating the swarm architecture and motion in real time, which is consistent with previous observations.(*16*) These results demonstrate the programmable nature of micrometre-sized Dynabot assemblies, enabling tuneable reconfiguration through external field parameters.

Building on this principle, we achieved controlled propulsion of Dynabot swarms in a vascular model of the human middle cerebral artery (MCA). Targeted navigation was accomplished by combining chain-walking locomotion induced by rotating magnetic fields with directional steering via superimposed magnetic field gradients (Fig. 5d-e and Movie S2). This hybrid control approach allowed precise positioning of the swarms within branched vascular geometries.

However, active propulsion against high physiological flow rates in large blood vessels remains challenging for microscale robotic agents. Alternatively, we explored swarm navigation within the cerebrospinal fluid (CSF) environment, where flow velocities are significantly lower (∼2.5 cm·s^-1^),(*42*) making it a more viable route for microcarrier-based drug delivery. This application holds particular promise for treating otherwise inoperable or diffuse neurological conditions, such as diffuse intrinsic pontine glioma (DIPG). To simulate this scenario, we deployed Dynabot swarms in a life-sized human ventricular phantom model and evaluated their navigability under clinically relevant conditions. Actuation using a rotating magnetic field (30 mT, 2.0 Hz) successfully propelled the swarms from the injection site at the median aperture, through the fourth ventricle and cerebral aqueduct, and into the third ventricle (Fig. 5f-g and Movie S3). The swarm exhibited stable, controllable locomotion throughout the curved anatomical pathways, confirming the feasibility of magnetically controlled swarm-based delivery in the central nervous system.

### In vitro navigation of monolithic Dynabots

To enhance *in vivo* traceability and mitigate unintended carrier loss during navigation, we fabricated larger-sized, monolithic Dynabots using a droplet evaporation strategy on a superamphiphobic surface, as schematically illustrated in Fig. 6a.(*43*) This top-down assembly approach yielded uniform spherical structures of hundreds of micrometers with pre-functionalized nanoparticles incorporated. SEM imaging and EDX elemental mapping confirmed the homogeneous distribution of zinc-ferrite, UiO-66-NH_2_, and tantalum nanoparticles within the spherical assemblies (Fig. 6b). Magnetic characterization of the monolithic Dynabots via superconducting quantum interference device (SQUID) magnetometry revealed a saturation magnetization of 78 emu g^-1^ for Dynabots with pure-ZFO, and 60 emu g^-1^ for the ones with all three components (Fig. 6c). Finally, the visibility of monolithic Dynabots under X-ray fluoroscopy was analyzed *ex vivo*, which confirmed the necessity of using composite-based Dynabots for reliable tracking through an ovine head (fig. S21).

**Fig. 6.**
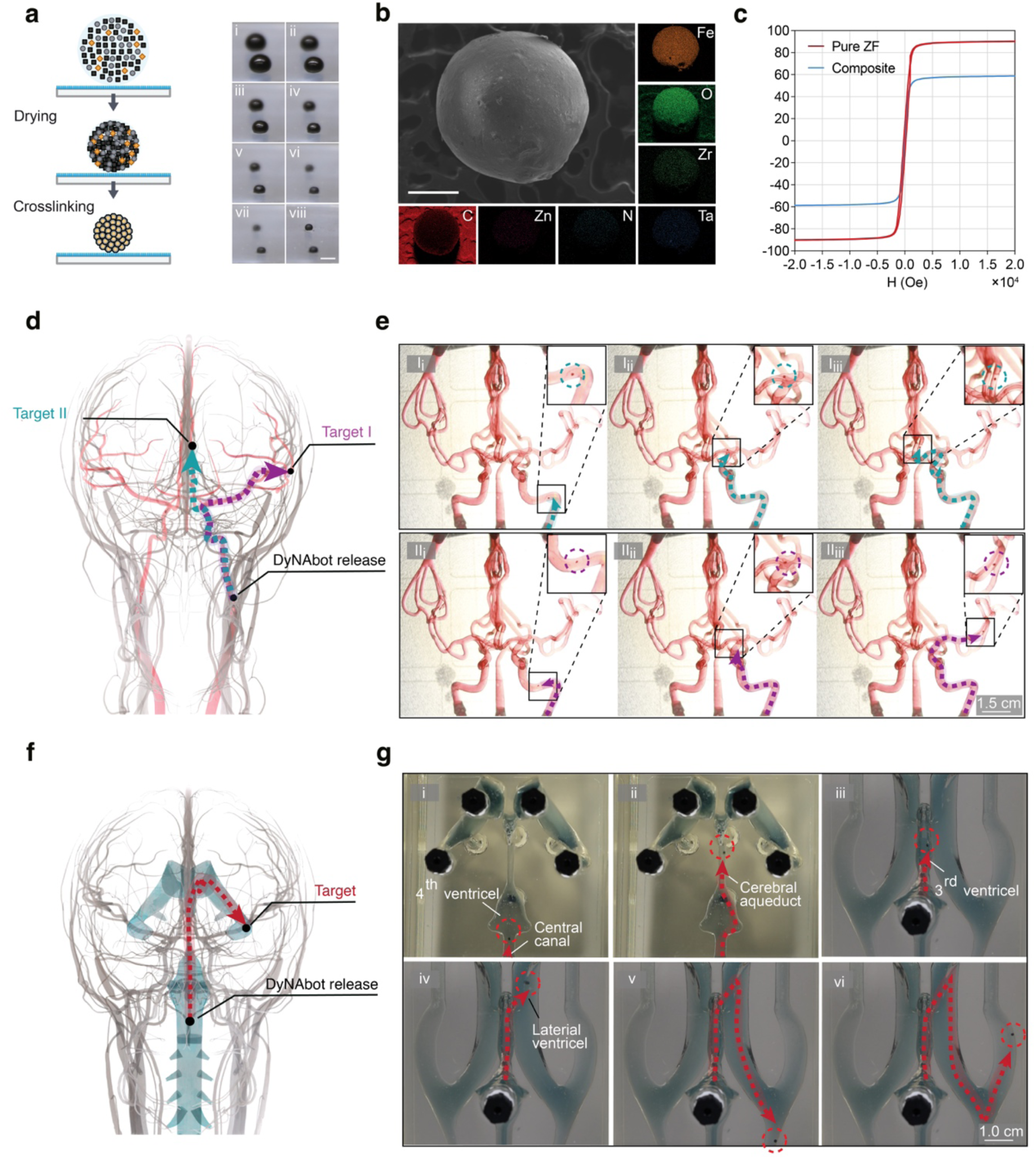
Monolithic Dynabot formation and *in vitro* navigation: **a**, Conceptual illustration and time-lapse images of macro-sized Dynabot formation on a superamphiphobic surface (scale bar: 2.5 mm). **b,** SEM images and EDX maps of macro-sized Dynabot (scale bar: 400 μm). **c,** Magnetic hysteresis loops at 300 K of macro sized Dynabots built from different nanoparticles (Composite) and only zinc-substituted iron oxide nanoparticles (ZF). **d,** Conceptual illustration of human head vasculature with highlighted area of phantom model. **e,** Time-lapse images of in-flow Dynabot navigation through a human head vasculature phantom model. **f,** Conceptual illustration of human head ventricular system, with the region corresponding to the phantom highlighted. **g,** Time-lapse images of Dynabot maneuvering through a human ventricular model.

Next, we assessed the navigation capability of the monolithic Dynabots under physiologically relevant flow conditions in a vascular phantom replicating the human internal carotid artery (ICA). Dynabots were injected into the ICA channel and subjected to a continuous flow of 35 cm·s^-1^, approximating arterial blood velocity in adult humans.(*44*) Upon applying a uniform magnetic field (30 mT) in combination with spatially varying magnetic gradients (350 mT·m^-1^), the monolithic Dynabots could be robustly steered toward pre-defined distal vessel targets (Fig. 6d-e and Movie S4). Moreover, their high magnetic responsiveness allowed for stable magnetic trapping within the ICA under flow, successfully overcoming hydrodynamic drag forces and keeping their positions at the target sites (Movie S5). These results confirm that monolithic Dynabots can function as magnetically navigable, clinically trackable carriers for targeted intravascular delivery.

Targeted delivery was furthermore demonstrated within a phantom model of the human ventricular system. A monolithic Dynabot was therefore released within the central canal and subsequently guided via the application of a magnetic field of 30 mT and varying gradients from the 4^th^ ventricle through the cerebral aqueduct towards the 3^rd^ ventricle, before being steered into the distal area of the lateral ventricle (Fig. 6f-g and Movie S6).

### In vivo demonstration of monolithic Dynabot actuation

To validate the potential clinical applicability of monolithic Dynabots, we conducted *in vivo* demonstrations in a porcine model under clinically relevant conditions. In the first experiment, a Dynabot was introduced into the common carotid artery (CCA) of the specimen, and a magnetic field of 30 mT combined with a gradient of 350 mT·m^-1^ was applied dorsally, perpendicular to the vessel axis. Following release, the Dynabot underwent flow-assisted magnetic targeting toward the ethmoidal artery (EA), consistent with the combined action of hydrodynamic drag and the externally applied magnetic force (Fig. 7a).

**Fig. 7.**
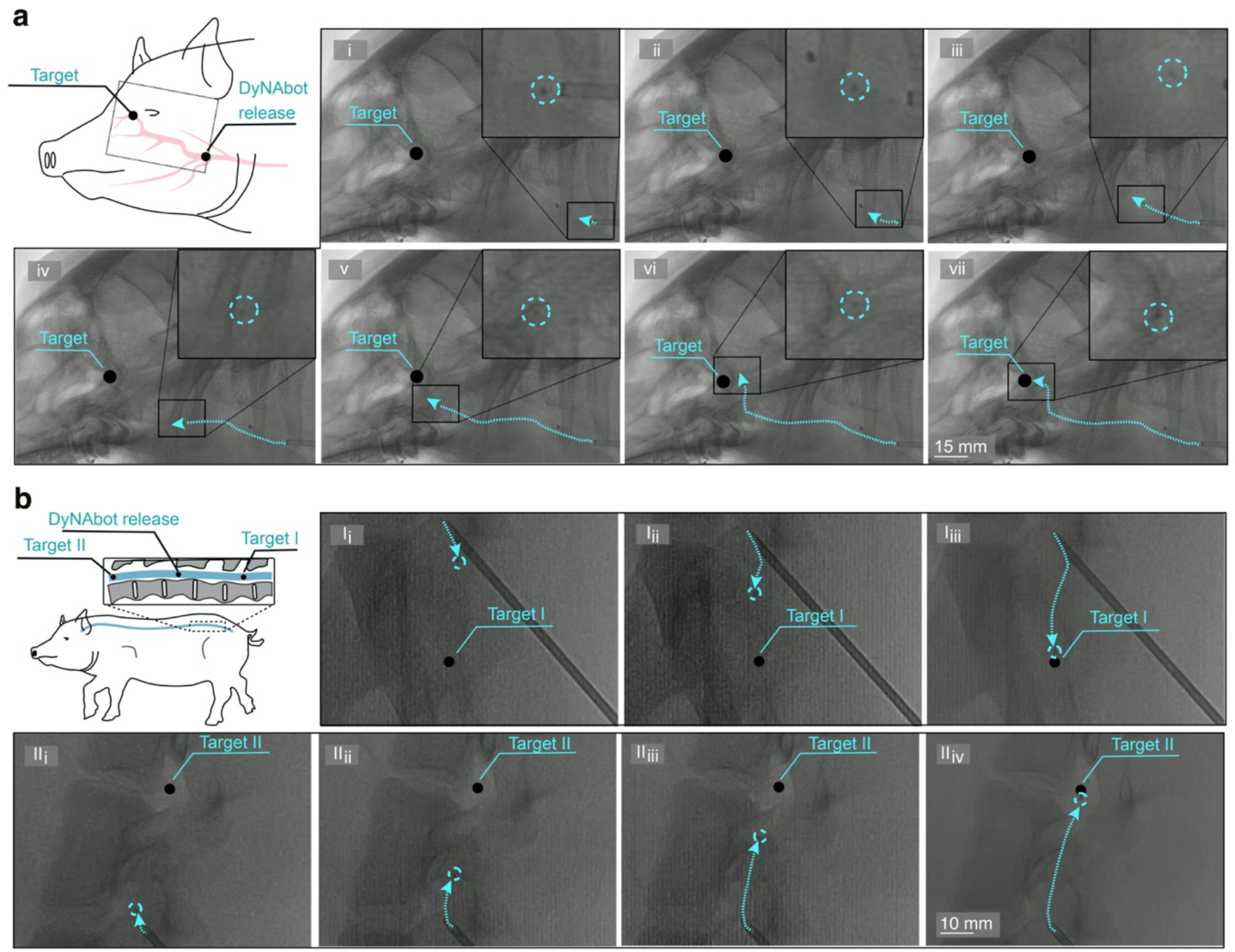
In vivo maneuvering of monolithic Dynabots within a clinical setting. **a**, Dynabot in-flow navigation from the common carotid artery (CCA) into the ethmoidal artery (EA). **b,** Dynabot navigation via rotating magnetic fields within the subarachnoid space.

To further assess the robots’ navigation capability within intricately structured subarachnoid spaces, two Dynabots were tested in the lumbar cistern, respectively. Using a 30 mT rotating magnetic field, the robots were guided bi-directionally along the cerebrospinal axis—toward both cranial and caudal directions—mimicking clinically relevant navigation pathways for neurotherapeutic delivery (Fig. 7b and Movie S7). These *in vivo* studies demonstrate that monolithic Dynabots are capable of being used in different anatomical compartments (including vascular and cerebrospinal environments) under dynamic physiological conditions. Their maneuverability and responsiveness *in situ* highlight their strong potential as versatile platforms for targeted and image-guided drug delivery in clinical applications.

## Discussion

In this study, we report the development of a dynamic, modular microrobotic platform, Dynabots, which are constructed from multifunctional nanoparticles assembled via thermally cleavable linkers. This system offers programmable structural transitions between assemblies and dispersed nanocarriers in response to thermal stimuli, enabling a seamless integration of magnetic navigation, controlled drug release, and imaging visibility. We demonstrate the robust synthesis and functional integration of magnetic, imaging, and drug-loading functionalities with nanoparticles as building blocks, followed by their assembly into programmable microscale and mesoscale architectures. The Dynabots exhibit excellent *in vitro* biocompatibility, enhanced drug delivery efficacy upon stimulus-induced disassembly, and potent antitumor performance both *in vitro* and *in vivo*. Furthermore, we validate their precise navigation capabilities within complex biological environments, including cerebral vasculature and ventricular systems, using both phantom models and live large-animal studies under clinically relevant conditions. These findings establish Dynabots as a highly adaptable microrobotic delivery platform that bridges the gap between precise magnetic control and responsive nanomedicine. Looking ahead, the modularity and programmability of Dynabots open up exciting possibilities for customized therapeutic strategies, multi-drug co-delivery, and integration with diagnostic functionalities. Future research may further explore autonomous control algorithms and long-term biocompatibility, advancing Dynabots toward clinical translation for minimally invasive, targeted treatment of complex diseases.

## Acknowledgements

1. L. H. and H. Y. contributed equally to this work. The authors thank the Nanoparticle Systems Engineering Laboratory (ETH Zurich, Prof. Dr Inge Herrmann) and the Medical Microsystems Laboratory (ETH Zurich, Prof. Dr Simone Schürle-Finke) for the usage of various characterization equipment. This work is supported by the National Natural Science Foundation of China (Project No. 52473254), Swiss National Science Foundation (Project No. 198643, and 190451), Grant PID2023–146623NB-I00 funded by MICIU/AEI/10.13039/501100011033/

FEDER/UE, Generalitat de Catalunya AGAUR grant 2021-SGR-00343, Maria de Maeztu Units of Excellence Programme CEX2023–001300-M and CEX2021-001202-M/funded by MCIN/AEI/10.13039/501100011033. A.L.O. acknowledges financial support from the grant CNS2022-135787 funded by MCIN/AEI/10.13039/501100011033 and European Union Next Generation EU/PRTR. J.P.-L. acknowledges financial support from Departament de Recerca i Universitats: del Departament d’Acció Climàtica, Alimentació i Agenda Rural; i del Fons Climàtic de la Generalitat de Catalunya (2023 CLIMA 00011) and the Generalitat de Catalunya (2021 SGR 00270).

## Competing interests

The authors declare no competing interests.

## Notes

### Competing Interest Statement

The authors have declared no competing interest.

