## Supporting Information for "Dynamic Nanoparticle Assembly-Based Biomedical Microrobots"

*^5^*International Institute for Intelligent Nanorobots and Nanosystems, College of Intelligent Robotics and Advanced Manufacturing, State Key Laboratory of Photovoltaic Science and Technology, Shanghai Frontiers Science Research Base of Intelligent Optoelectronics and Perception, Institute of Optoelectronics, Fudan University, Shanghai, 200433, China. *^6^*Zhejiang Key Laboratory of Extreme Environment Functional Materials, Yiwu Research Institute of Fudan University, Yiwu, 322000, China

*^7^*Departament de Física, Universitat Politècnica de Catalunya, BarcelonaTech (UPC). Institut de Tècniques Energètiques (INTE). Barcelona Research Center in Multiscale Science and Enigneering, Av. Eduard Maristany 16, 08019 Barcelona, Spain

*^8^*Departamento de Ciencias, Universidad Pública de Navarra, Campus de Arrosadía, 31006, Pamplona, Spain. Institute for Advanced Materials and Mathematics (INAMAT^2^), Universidad Pública de Navarra, Pamplona E-31006, Spain

*^9^*Departament de Ciència dels Materials i Química Física Institut de Química Teòrica i Computacional, University of Barcelona, Barcelona 08028, Spain

*^10^*Institució Catalana de Recerca i Estudis Avançats (ICREA), Pg. Lluís Companys 23, 08010 Barcelona, Spain

*^11^*Center for Preclinical Development, University Hospital Zurich, University of Zurich, Zurich, Switzerland

**Experimental Section**

**Synthesis of Nitrodopamine**

Nitrodopamine hydrogensulfate was synthesized following a previously established protocol. In a typical synthesis, dopamine hydrochloride (Sigma-Aldrich, CAS: 62-31-7) (500 mg, 2.625 mmol) was first dissolved in 15 mL of deionized water before adding sodium nitrite (Sigma-Aldrich, CAS: 7632-00-1) (624.5 mg, 9.05 mmol) and cooling the mixture to 0 °C. Next, 5 mL of sulfuric acid (20% v/v) (Sigma-Aldrich, CAS: 7664-93-9) was added dropwise to the mixture under vigorous stirring, while maintaining the reaction flask in an ice bath. During the addition, the mixture changed color from translucent to muddy brown, followed by the formation of a yellow precipitate, indicating the formation of nitrodopamine hydrogensulfate. After the addition, the mixture was allowed to stir at room temperature overnight, after which the crude product was filtered and washed five times with ice-cold deionized water. Lastly, the collected nitrodopamine hydrogensulfate was dried under high vacuum and stored at 4 °C for further use.

**Magnetic nanoparticle synthesis**

Cubic-shaped zinc-substituted iron oxide nanoparticles were synthesized via thermal decomposition by adjusting the previously introduced synthesis methods.^25^ In a typical synthesis, Fe(acac)_3_ (Thermo Fischer, CAS: 14024-18-1) (424 mg, 1.2 mmol), Zn(acac)_2_ (Sigma-Aldrich, CAS: 14024-63-6) (79.11 mg, 0.3 mmol), and sodium oleate (TCI, CAS: 143-19-1) (200 mg, 0.657 mmol) were dissolved in benzyl ether (Thermo-Scientific, CAS: 103-50-4), which had been oxidized for 17 hours at 60 °C prior to mixing. Subsequently, 1-octadecene (Sigma-Aldrich, CAS: 112-88-9) (15 mL), 1-tetradecene (Sigma-Aldrich, CAS: 1120-36-1) (3 mL), and oleic acid (OA, Sigma-Aldrich, CAS: 112-80-1) (1.6669 g, 5.907 mmol) were added, and the mixture was homogenized via sonication. The resulting slurry was then degassed at 60 °C for 60 minutes under vigorous stirring before being heated under N_2_ flow to 294 °C with a heating rate of 3 °C min^-1^. The mixture was kept at reflux for 90 minutes. The resulting crude product was allowed to cool to room temperature before being stored at 4 °C for further processing.

**Particle purification**

The as-synthesized oleic acid-coated zinc-substituted iron oxide nanoparticles were purified by a sequence of dispersion and sedimentation steps in a mixture of different solvents, as previously described.^25^ In a typical process, 80 mg of the as-synthesized nanoparticles were first washed twice in a mixture of chloroform (Merck, CAS: 67-66-3) (25 mL) and acetone (Sigma-Aldrich, CAS: 67-64-1) (75 mL) by particle precipitation via centrifugation, followed by re-dispersion in chloroform. Next, the particles were washed twice more by re-dispersion in chloroform (25 mL) and precipitation in a mixture of methanol (Sigma-Aldrich, CAS: 67-56-1) (50 mL) and acetone (50 mL) via centrifugation. Finally, the purified oleic acid-coated nanoparticles were dispersed in dry DMF (Acros Organics, CAS: 68-12-8) at a concentration of 10 mg mL⁻¹ for further processing.

**UiO-66 NH_2_ nanoparticle synthesis**

UiO-66 NH_2_ nanoparticles were synthesized by dissolving zirconyl chloride octahydrate (Sigma-Aldrich, CAS: 13520-92-8) (9.12 mg, 0.028 mmol) in dry DMF (0.5 mL), and 2-aminoterephthalic acid (Sigma-Aldrich, CAS: 10312-55-7) (5.13 mg, 0.028 mmol) in dry DMF via sonication. Subsequently, the two solutions were added to 0.769 mL of dry DMF and allowed to react at 120 °C for 2 hours. Afterwards the particles were washed four time by centrifugation, followed by re-disperion in DMF.

**Ligand modifications of nanoparticles**

Functionalization of the different types of nanoparticles (as-synthesized ZF nanoparticles, UiO-66 NH_2_ nanoparticles, and Tantalum nanoparticles (Sigma-Aldrich, CAS: 7440-25-7)) with nitrodopamine was carried out according to a previously established surface modification protocol.^25^ In a typical reaction, 40 mg of nanoparticles were dispersed in dry DMF (Acros Organics, CAS: 68-12-8) (4 mL), while constantly bubbling N_2_ through the mixture. Subsequently, 13.3 mg of synthesized nitrodopamine hydrogensulfate was added to the dispersion. After 10 minutes of N_2_ purging, the mixture was sonicated for 60 minutes, before being allowed to react at room temperature for 24 hours. Afterward, the functionalized nanoparticle-containing mixture was sonicated once more for 60 minutes before precipitating the nitrodopamine-functionalized nanoparticles in 100 mL of cold acetone. Subsequently, the particles were washed five more times by centrifugation and re-dispersion in methanol, before being transferred into dry DMF, where they were washed two more times by re-dispersion and centrifugation, and then re-dispersed in a concentration of 10 mg/mL.

**Diels-Alder Precursor Synthesis**

The synthesis of the Diels-Alder precursor followed a previously established method with slight adjustments.^13^ In a clean amber vial, 3-(2-furyl) propionic acid (Sigma-Aldrich, CAS: 935-13-7) (168 mg, 1.2 mmol) and 3-maleimidopropionic acid (ThermoFisher Scientific, CAS: 7423-55-4) (203 mg, 1.2 mmol) were dissolved in 1 mL of a 1:1 mixture of DMF and DI water under a N_2_ atmosphere by sonication for 5 minutes. Following this, the solution was stirred at 40 °C for 72 hours on a heating plate, before being cooled to room temperature and the solvents were removed under high vacuum. The resulting crude product was then precipitated in chloroform and dried via vacuum filtration, resulting in the collection of a white powder product.

**ThermoXlinker Synthesis**

The thermo-responsive crosslinking agent (ThermoXlinker) was synthesized according to a previously established protocol.^13^ In a typical reaction, the previously synthesized DA precursor (74 mg, 0.24 mmol) was dissolved in dry DMF (1 mL) inside a N_2_-filled glovebox. To this, HBTU (Sigma-Aldrich, CAS: 94790-37-1) (227 mg, 0.6 mmol), DIPEA (Sigma-Aldrich, CAS: 7087-68-5) (108 mg, 0.84 mmol), and Azide-PEG3-Amine (Broadpharm, CAS: 134179-38-7) (110 mg, 0.6 mmol) were added. The solution was then stirred at room temperature for 16 hours in the dark. After the reaction was complete, the solvent was removed under high vacuum, and the resulting highly viscous crude product was dissolved in DCM (Sigma-Aldrich, CAS: 75-09-2) (1 mL). The product was subsequently purified by column chromatography over silica gel, starting with a 25:1 DCM:methanol mixture as eluent and gradually increasing to 15:1, yielding a highly viscous yellow product.

**THPTA-Cu complex**

The THPTA-Cu complex for copper-catalyzed azide-alkyne cycloaddition was prepared by mixing a 50 mM THPTA (TCI, CAS: 760952-88-3) (500 µL) solution with a 20 mM copper(II) sulfate (Sigma-Aldrich, CAS: 7758-98-7) (250 µL) solution.

**Nanoparticle Crosslinking**

Crosslinking of the alkyne-functionalized nanoparticles was carried out by dispersing 50 mg of nanoparticles (weight ratio of 70:25:5 ZF:Ta:UiO66-NH_2_) and 5 mg of the synthesized ThermoXlinker in 5 mL of de-oxygenated DI water, followed by the addition of freshly prepared THPTA-Cu complex (50 µL) and sodium ascorbate (Sigma-Aldrich, CAS: 134-03-2) (50 µL, 25 mg/mL). Crosslinking of DBCO-functionalized nanoparticles was performed via the SPAAC reaction by adding 5 mg of the synthesized ThermoXlinker to a 10 mg/mL dispersion of DBCO-terminated nanoparticles (weight ratio of 70:25:5 ZF:Ta:UiO66-NH_2_) in 5 mL of solution.

**Microscopy**

*Scanning Electron Microscopy (SEM):* SEM images were recorded using a Zeiss ULTRA 66 (Carl Zeiss GmbH, Oberkochen, Germany) operating at 5 kV. EDX mapping analysis was performed with a 60 µm aperture and a 10 mm working distance at 20 kV.

*Transmission electron Microscopy (TEM):* TEM, scanning TEM, and EDX analysis of nanoparticles were performed using an FEI Talos F200X (Chem S/TEM) (Thermo Fisher Scientific Inc., Waltham, Massachusetts, USA) operating at 200 kV. Samples were prepared by drop-casting diluted particle dispersions onto a carbon-coated Cu TEM grid (400 mesh) (1824, Ted Pella).

**Structural analysis**

*Fourier Transform Microscopy (FTIR):* Spectra of the samples were recorded using a Varian 640 Fourier Transform Infrared Spectrometer equipped with a Golden Gate diamond ATR (Varian Inc., Massachusetts, USA) across a range of 4000–400 cm⁻¹ with a resolution of 4 cm⁻¹.

*Nuclear Magnetic Resonance (NMR) Spectroscopy:*¹H-NMR spectra were collected using a 400 MHz Bruker Avance Ultrashield (Bruker Corp., Billerica, Massachusetts, USA), with DMSO-d₆ as the solvent.

*Thermogravimetric Analysis (TGA*): Thermogravimetric measurements of dry samples were performed using a Mettler Toledo TGA/DSC 3+ Star (Mettler-Toledo, Columbus, Ohio, USA) under a steady O₂ flow of 60 mL min⁻¹, heating from 30 °C to 900 °C at a rate of 10 °C min⁻¹.

*X-ray Diffraction (XRD) analysis:* The crystal structure of nanoparticle powders was determined using a Malvern Panalytical Empyrean diffractometer (Malvern Panalytical GmbH, Kassel, Germany) equipped with a copper X-ray source (λ = 1.5406 Å) and a PIXcel detector. Measurements were performed in the range of 4°≤ 2θ ≤ 80°, with a step size of 0.04° and a sweep rate of 0.2 seconds per step. Crystallite sizes were calculated from the main Bragg peak with the highest intensity using the Scherrer equation.

*Small Angle X-ray Scattering (SAXS):* SAXS measurements were performed on a Bruker AXS Micro instrument, with a microfocused X-ray source, operating at voltage and filament current of 50 kV and 1000 μA, respectively. The Cu Kα radiation (λ CuKα = 1.5418 Å) was collimated by a 2D Kratky collimator, and the data were collected by a 2D Pilatus 100K detector. The scattering vector q = (4π/λ) sin θ, with 2θ being the scattering angle, was calibrated using silver behenate. Data were collected and azimuthally averaged using the Saxsgui software to yield 1D intensity vs. scattering vector q, with a q range from 0.004 to 0.5 Å^–1^. For all measurements the samples were placed inside a stainless-steel cell between two thin replaceable mica sheets and sealed by an O-ring, with a sample volume of 10 μL and a thickness of ∼1 mm. Measurements were performed at 20°C, and samples were equilibrated for 15 min before measurements, whereas scattered intensity was collected over 20 min.

*Dynamic light scattering (DLS) & ζ Potential measurements:* Hydrodynamic diameters and zeta potentials of diluted particle dispersions in de-ionized water and PBS were measured using a Malvern Anton Paar Litesizer 500 DLS (Anton Paar Group AG, Graz, Austria) at 32 °C.

*Transmission Mössbauer Spectroscopy:* Transmission Mössbauer Spectra (TMS) were obtained in transmission mode at room temperature and pressure using a conventional spectrometer with constant acceleration with a 25 mCi ^57^Co radioactive source in Rh matrix. The spectra were recorded in a standard multichannel analyzer using a velocity range of ± 12.2 mm sec^-1^ and were subsequently fitted with the NORMOS software. Sample fitting was undertaken with one or several distributions of hyperfine and one paramagnetic singlet. The fitted parameters are the isomer shift (δ), always expressed relative to the isomer shift of the bcc-Fe at room temperature, the quadrupole splitting (Δ), and the hyperfine magnetic field (BHF). In all cases, the area expressed in % corresponds to the fraction of Fe atoms in a particular environment with respect to the total amount of Fe atoms, and the magnitudes inside parentheses are the standard deviation.

*Mössbauer spectroscopy:* Mössbauer spectra were measured in transmission mode at 300 K using a conventional spectrometer that was equipped with a 25 mCi 57 Co radioactive source in a Rh matrix. Spectra were recorded with constant acceleration on a standard multichannel analyzer using a velocity range of ± 12.2 mm s^-1^, and subsequently fitted with the NORMOS software. Fitting was undertaken by using several hyperfine field distributions and one paramagnetic singlet.

**Magnetic characterization**

*Vibrating samples magnetometry:* Symmetric magnetic hysteresis loops were measured for nanoparticles and capsules over the field range 2 T at 300 K, using a vibrating sample magnetometer (VSM EZ9, Microsense). Nanoparticle samples were prepared by drop casting highly concentrated dispersions of oleic acid respectively nitro dopamine functionalized particles onto a round filter-paper substrate (Ø: 8mm) and drying under a high vacuum for 48 hours. Normalization to magnetization per gram metal-ferrite was undertaken by accounting for organic weight percentage loss determined *via* TGA.

*AC-Magnetometer:* The dynamic magnetization of the nanoparticles under alternating magnetic field was measured using a lab-made AC Magnetometer equipped with a fiber-optic temperature sensor (Neoptix).^43^ The specific absorption rate (SAR) was calculated from the area of the M(H) loops using the following equation:

$$SAR\text{=-}f\oint_{cycle} M_{t}{dH}_{app}$$

where *f* and *H*_app_ are the frequency and intensity of the externally applied magnetic field and *M*_t_ is the dynamic magnetization of the nanoparticles (in mass magnetization units).

*AC hysteresis loop and SAR numerical simulations:* The numerical simulations were performed solving the Landau–Lifshitz–Gilbert (LLG) equation as described in [DOI: https://doi.org/10.1103/PhysRevE.90.023203]. The particle volume and elongation were inferred from TEM images (17.9x17.9x25.6 nm, see Figure 2a), from which a magnetic shape anisotropy of 14 kJ.m^-3^ was inferred. For magnetite, an intrinsic saturation magnetization of 400 kA.m^-1^ was used whereas a temperature of 300 K was assumed. When accounting for interparticle interactions, the following dipolar effective field was considered:

$$H_{dip,i}\text{=}\frac{1}{4\pi}\sum_{j\neq i} 3\frac{\left( m_{j}\cdot\hat{r}_{ij}\text{-}m_{j} \right)}{r_{ij}^{3}}$$

where, $m_{j}$ is the magnetic moment of each particle and $r_{ij}$ is the interparticle distance between particles in the assembled Dynabot. The position, and therefore the interparticle distances, were assigned using the Gmsh mesh generator, enforcing an interparticle distance of 41 nm (26/2+15 nm), being each node the position of each particle.

Each simulation was carried out with 4452 individual IO particles and the error was estimated from standard deviation. The rotational motion of each particle was neglected.

*Superconducting Quantum Interference Device:* The magnetic properties of dried sample powders were measured in a compacted state using a PPMS-9 (Quantum Design), equipped with a cryostat that can measure from 2 to 400 K, between -7 and 7 Tesla. Magnetic hysteresis curves were recorded at 300K and 5K.

**Cell culture**

Mouse NIH/3T3 fibroblasts and B16-F10 murine melanoma cells were obtained from the American Type Culture Collection (ATCC) and cultured according to standard protocols. Both cell types were maintained in high-glucose Dulbecco’s modified Eagle’s medium (Gibco, 10566016), supplemented with 10% (v/v) fetal bovine serum (Gibco, A5670701), penicillin (100 units ml^-1^), and streptomycin (100 μg ml^-1^). Cells were incubated at 37  °C in a humidified atmosphere containing 5% CO_2_. Subculturing was performed using 0.25% trypsin-EDTA (Gibco, 25200056) in T-75 culture flasks.

**Biocompatibility**

Biocompatibility of the samples was evaluated using NIH/3T3 or B16-F10 cells. Samples were thoroughly rinsed with phosphate-buffered saline (PBS) and sterilized by ultraviolet (UV) irradiation overnight. NIH/3T3 fibroblasts were seeded in 96-well plates at a density of 4 × 10^3^ cells per well. After 24 h of attachment, cells were exposed to the sterilized samples and incubated for 72 h under standard culture conditions. Cell viability was quantified via the MTT assay (3-(4,5-dimethylthiazol-2-yl)-2,5-diphenyltetrazolium bromide) at a final concentration of 3 mg/mL. Absorbance was measured at 570 nm with a reference wavelength of 630 nm using a Tecan multimode microplate reader (Infinite 200 PRO). In parallel, live/dead staining was performed using a membrane-permeable calcein-AM and membrane-impermeable ethidium homodimer-1 solution (Live/Dead Viability/Cytotoxicity Kit, Invitrogen, Cat. No. R37601). After 15 min incubation at room temperature, cells were imaged using a Zeiss LSM 880 confocal microscope equipped with Airyscan. Image analysis and quantification were conducted using ImageJ software.

**Cellular uptake and cytotoxicity assays**

To evaluate the therapeutic potential and biocompatibility of Dynabots, we first loaded the assembled structures with doxorubicin (DOX) by passive absorption. Cellular uptake was then investigated under magnetic hyperthermia (MHT) and ultrasound stimulation, conditions that trigger Dynabot disassembly. For uptake studies, B16-F10 cells (1 × 10^5^ per well) were seeded in 12-well plates and cultured for 24 h, followed by incubation with DOX-loaded Dynabots (DOX concentration 2 μg mL^-1^) for 2 h. After three washes with ice-cold PBS, cell nuclei were counterstained with DAPI and examined using confocal laser scanning microscopy (LSM880, Zeiss). Quantification of intracellular DOX was performed by flow cytometry (LSRFortessa, BD Biosciences).

For cytotoxicity evaluation, B16-F10 cells (4 × 10^3^ per well) were seeded in 96-well plates and incubated for 24 h, then exposed to samples at varying concentrations for an additional 24 h. Cell viability was assessed using the MTT assay, and the absorbance was measured on a microplate reader.

**Tumour penetration in 3D spheroids**

Three-dimensional tumour spheroids were established using ultra-low attachment 384-well spheroid microplates (Corning, 4516). B16-F10 cells (1,000 cells per well) were seeded in complete medium supplemented with 10% fetal bovine serum and cultured at 37 °C until spheroids reached an average diameter of ~300 µm. Spheroid growth was monitored by optical microscopy, and only uniform, compact spheroids were selected for experiments. The tumor penetration was then investigated under MHT and ultrasound stimulation. For penetration studies, the B16-F10 spheroids were transferred to 35 mm confocal dishes and incubated with various samples for 4 h and then under MHT and ultrasound stimulation. Doxorubicin (DOX) fluorescence within Dynabots was imaged using a confocal laser scanning microscope by Z-stack tomography from the spheroid surface to the equatorial plane. The mean fluorescence intensity was quantified using ImageJ software.

**Mice**

Male C57BL/6 mice were maintained at the Animal Center of Shenyang Pharmaceutical University, China. All procedures were performed in accordance with protocols approved by the Institutional Animal Ethical Care Committee of the university (approval no. SYXK 2021-0007, Liaoning, China).

**Tumour models and therapy**

For the B16-F10 melanoma model, 1×10^6^ B16-F10 cells were injected subcutaneously into the right flank of C57BL/6 mice. When tumor volumes reached approximately 150 mm^3^, mice received peritumoral injections of Dynabots containing doxorubicin 1 mg kg^-1^ once daily for 6 consecutive days. After each injection, tumors were irradiated with 808 nm near-infrared light for 5 min at a power density of 1 W cm^-2^ and then followed with ultrasonic vibration using a hand-held ultrasound device (Roscoe Medical, US Pro 2000 2^nd^ edition) at low power (0.32 W ± 20 %) for 1 min. Body weight and tumor size were recorded every 2 d. On day 20, blood was collected for hepatic and renal function analyses, major organs were harvested for hematoxylin and eosin staining, and excised tumors were processed for penetration assessment by confocal microscopy.

**Finite-element thermal modelling**

Finite-element thermal modelling was performed using COMSOL Multiphysics to estimate the local temperature distribution around the peritumorally injected Dynabot-rich region during 808 nm NIR irradiation. The injected material was modeled as a dispersed disc-shaped heat-generating domain embedded in biological tissue. Effective density, heat capacity and thermal conductivity were estimated from the mass fractions of Zn-substituted iron oxide nanoparticles, tantalum nanoparticles, UiO-66-NH2 nanoparticles and ThermoXlinker. The laser input was represented as a volumetric heat source corresponding to the absorbed optical power. The radius and depth of the heat-generating region were swept to bracket plausible in vivo distributions. The tissue boundary was maintained at body temperature. Blood perfusion was neglected over the short irradiation window, and the resulting temperatures should therefore be interpreted as an upper-bound estimate.

**Swarm Formation**

Magnetic field dependent swarm formation was assessed in a custom-built electromagnetic set-up varying the magnetic field strength from 10 – 40 mT, and its rotational frequency from 0-8 Hz.

**In vitro navigation**

All navigation tests were performed in water at room temperature. For actuation within the brain vasculature and ventricular phantom models red or blue food coloring was added for better visibility.

*Swarm manipulation:* Swarm maneuvering in three-dimensional spaces was demonstrated in custom-made, patient data-based vascular and ventricular silicone models (Swiss Vascular). Actuation was performed either by varying rotational field frequencies in a magnetic field of 30 mT, applied using an Octomag electromagnetic navigation system (Magnebotix), or at 30 mT and 2 Hz, within an electromagnetic navigation system consisting of two Navion sub-units (Magnebotix AG) facing each other.

*In vitro in flow navigation:* Three-dimensional navigation was demonstrated with monolithic Dynabots within a patient data-based, custom-made silicone vascular model of the human head vasculature (Swiss Vascular) following a previously established protocol.^18^ The fluid flow entering the internal carotid artery (ICA) and basilar artery (BA) was adjusted to 35 cm/s and 30 cm/s, using a flow sensor (SLF3S-400B, Sensirion). In a typical demonstration, a magnetic gradient was applied towards the direction of the the targeted vasculature branch (30 mT, 350 mT/m), using an electromagnetic navigation system (eMNS), comprising two Navion units (Magnebotix AG) placed 30 cm apart, which were positioned perpendicular to the model.

**In vivo electromagnetic actuation study**

The animal study was approved by the local Committee for Animal Experimental Research (Cantonal Veterinary Office Zurich, Switzerland) under license number ZH072/2023. The experiments were conducted on intact female pig, weighing approximately 50 kg. On the day of the procedure, the animal was sedated via intramuscular injection of ketamine (Ketasol®-100 ad us.vet.; Dr. E. Graeub AG, Berne, Switzerland) at 15 mg/kg body weight (BW), azaperone (Stresnil® ad us.vet.; Elanco Tiergesundheit AG, Basel, Switzerland) at 2 mg/kg BW, and atropine (Atropinsulfat KA 1%; Kantonsapotheke, Zurich, Switzerland) at 0.05 mg/kg BW. Anesthesia was induced with intravenous propofol (Propofol® Lipuro 1%; B. Braun Medical AG, Sempach, Switzerland) at 1–2 mg/kg BW via a vein in the auricle (V. auricularis), suppressing the swallowing reflex to allow intubation. Anesthesia was maintained with continuous IV infusion of propofol (Propofol® Lipuro 2%) at 3 mg/kg BW/h, along with inhaled isoflurane (Attane™ Isoflurane ad us.vet.; Piramal Enterprises, India) at 1.0–1.5% in a 70% oxygen-to-air mixture via mechanical ventilation. A 7F guide catheter was inserted through an 8F femoral access sheath and navigated under fluoroscopic guidance to one of the common carotid arteries (CCA). The monolithic Dynabot was delivered through the catheter and released into the bloodstream within the CCA. In some instances, Dynabots were also introduced into the subarachnoid space via lumbar puncture using a 14G needle. Fluoroscopic imaging was performed using a mono-planar system (Allura Xper FD20, Philips N.V.) at 30 frames per second. The electromagnetic navigation system (eMNS), comprising two Navion units (Magnebotix AG) placed 30 cm apart, was positioned perpendicular to the operating table.

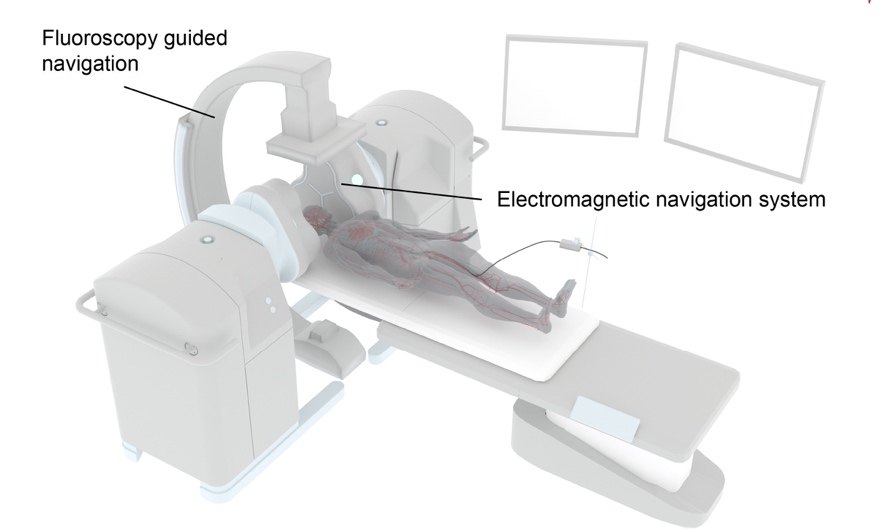

**Fig. S1.** Conceptional illustration of the utilized clinical set-up for microrobot maneuvering.

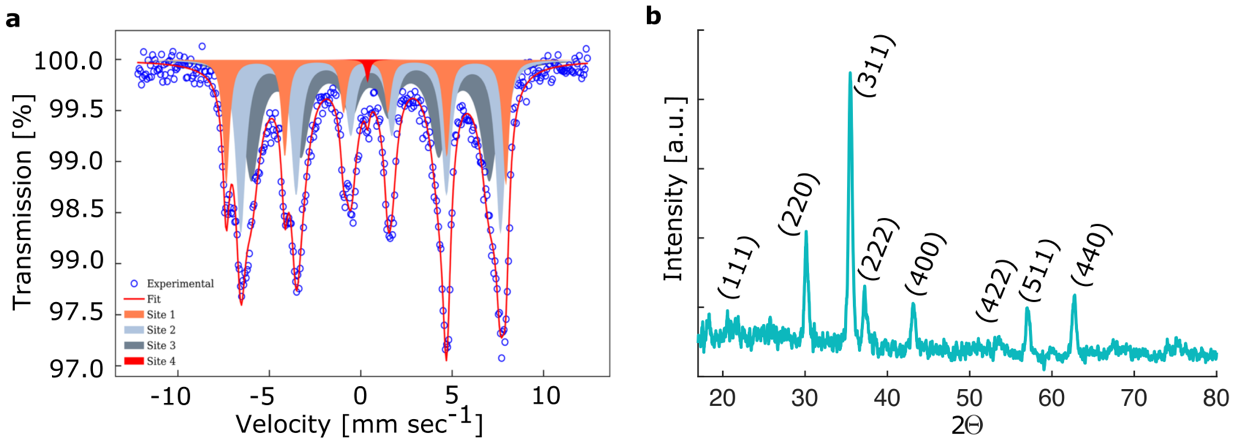

Fig. S2. (a) Mössbauer spectra of as synthesized zinc-substituted iron oxide nanoparticles. (b) XRD spectra of as synthesized zinc-substituted iron oxide nanoparticles

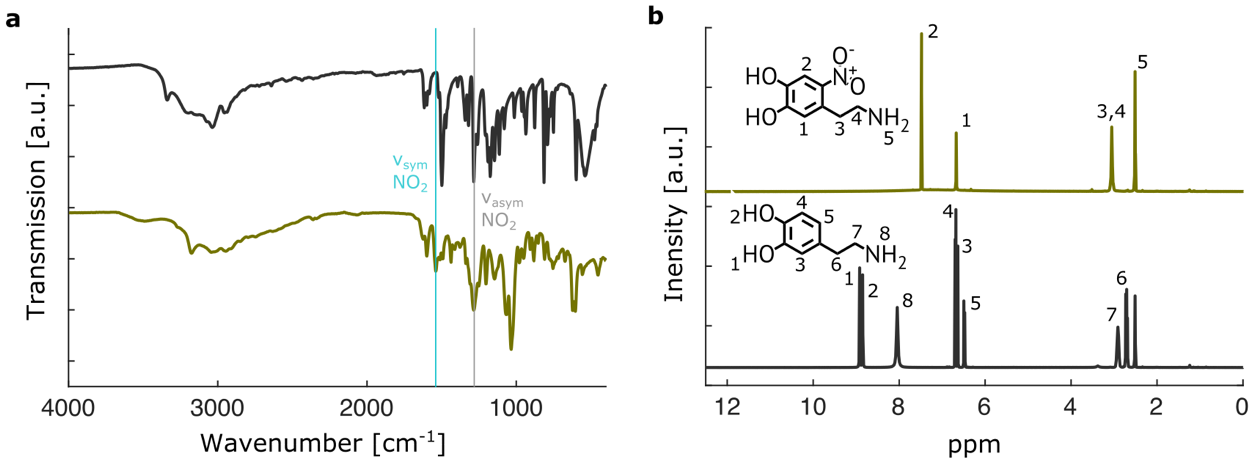

Fig. S3 Characterizations of nitrodopamine synthesis: (a) FTIR spectra of dopamine (black) and nitrodopamine hydrogen sulfate (yellow). (b) NMR spectra of dopamine (black) and nitrodopamine hydrogen sulfate (yellow).

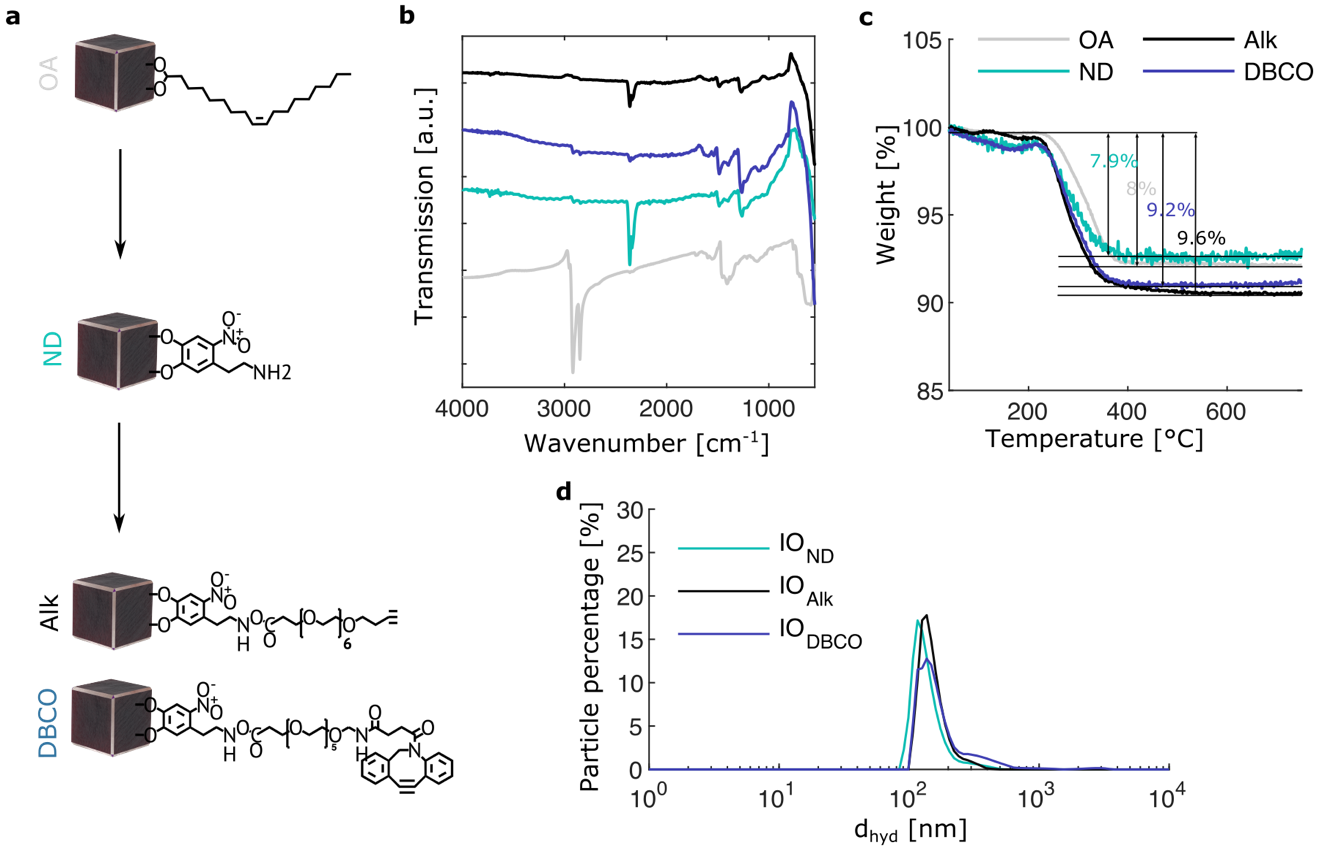

Fig. S4. (a) Schematic illustration of undertaken particle surface functionalizations of zinc-substituted iron oxide nanoparticles. (b) FTIR spectra of dried nanoparticle powders with different surface functionalizations. (c) TGA analysis of dried nanoparticle powders with different surface functionalizations. (d) DLS spectra of colloidal nanoparticle dispersions in DI-water of zinc-substituted iron oxide nanoparticles after the different surface functionalizations.

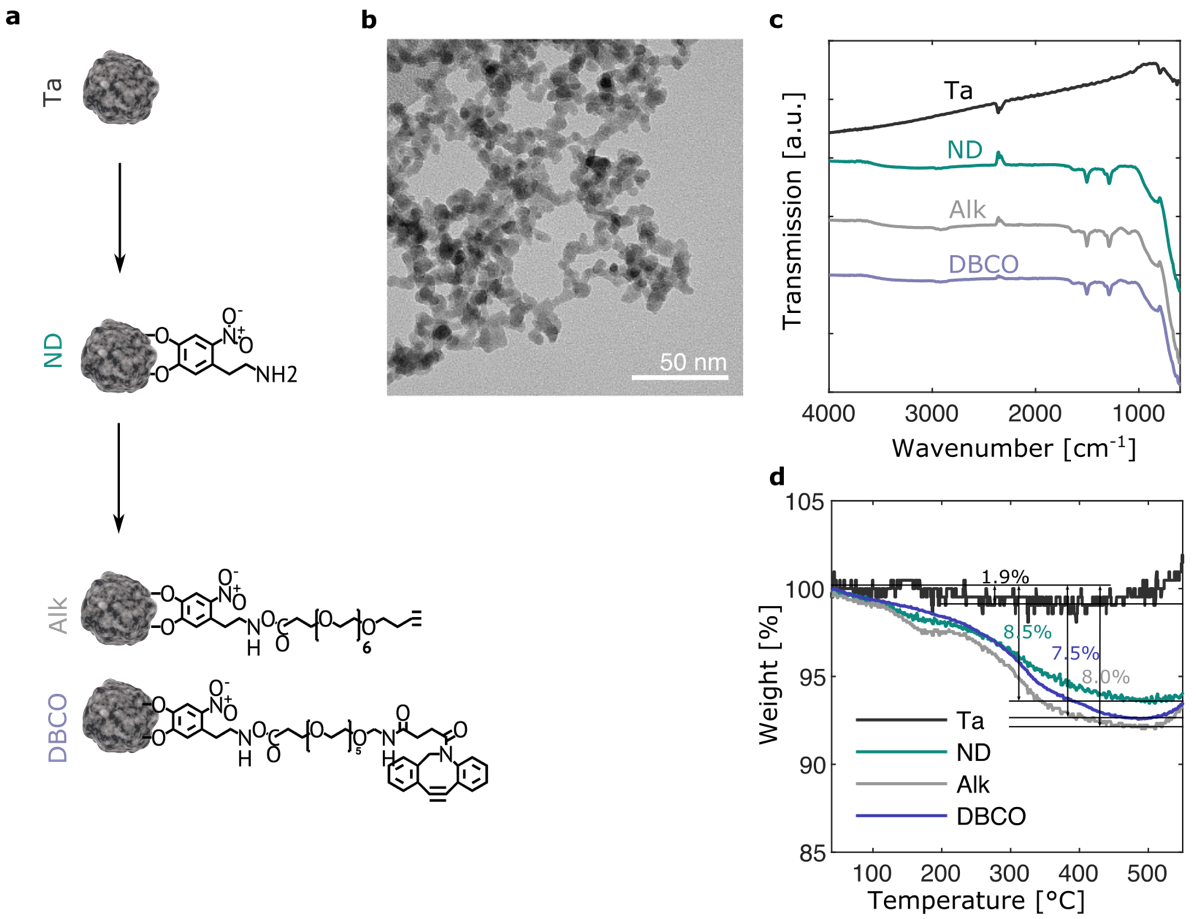

Fig. S5. (a) Schematic illustration of undertaken particle surface functionalizations of tantalum nanoparticles. (b) TEM image of tantalum nanoparticles (c) FTIR spectra of dried nanoparticle powders with different surface functionalizations. (d) TGA analysis of dried nanoparticle powders with different surface functionalizations.

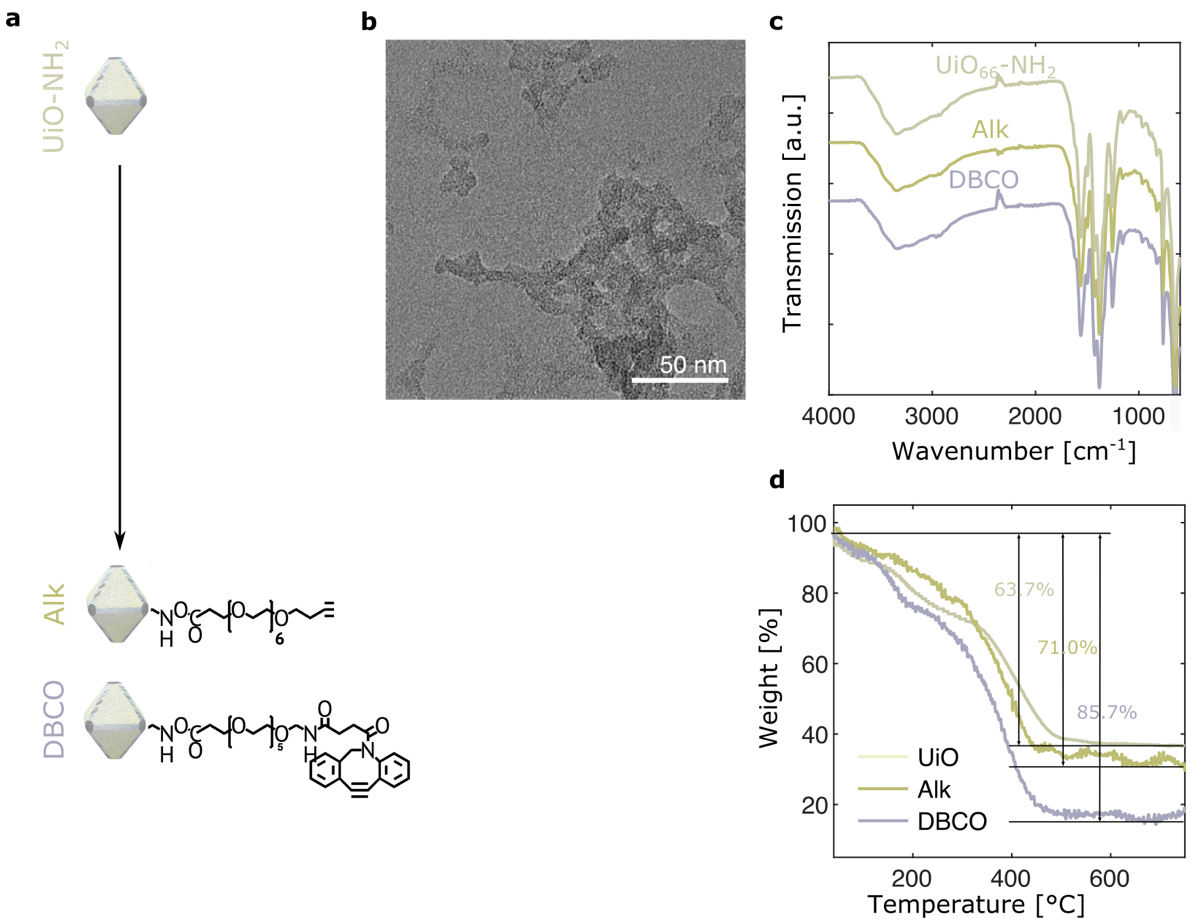

Fig. S6. (a) Schematic illustration of undertaken particle surface functionalizations of UiO_66_-NH_2_ nanoparticles. (b) TEM image of UiO_66_-NH_2_ nanoparticles (c) FTIR spectra of dried nanoparticle powders with different surface functionalizations. (d) TGA analysis of dried nanoparticle powders with different surface functionalizations.

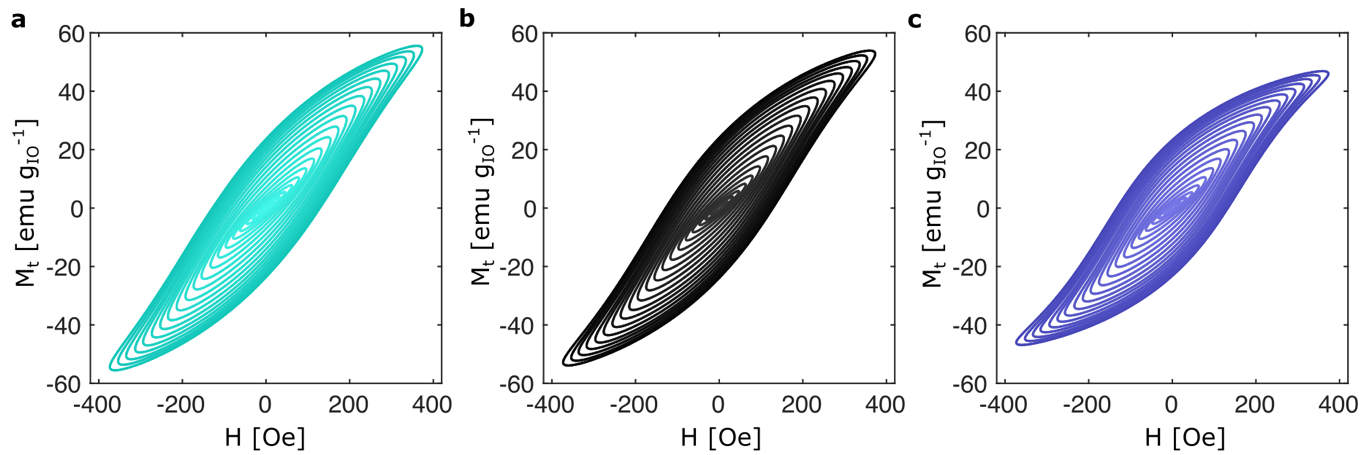
Fig. S7. Field strength dependent AC hysteresis loops of different functionalized Zn-substituted iron oxide nanoparticles. (a) ND. (b) Alk. (c) DBCO.

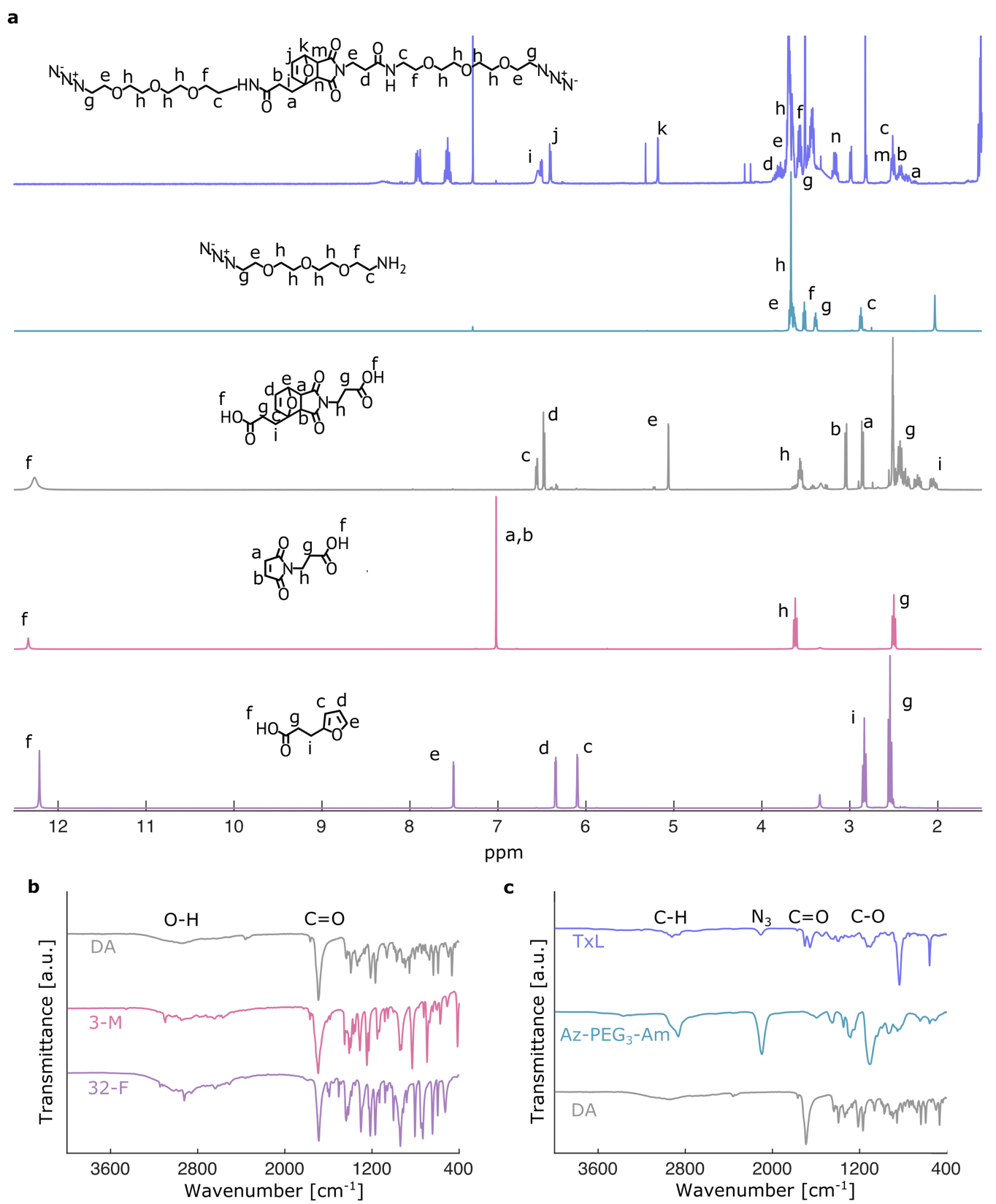

Fig. S8. (a) NMR spectra of the ThermoXlinker and its reagents. (b) FTIR spectra of the DA adduct and its reagents. (c) FTIR spectra of the ThermoXlinker and its reagents

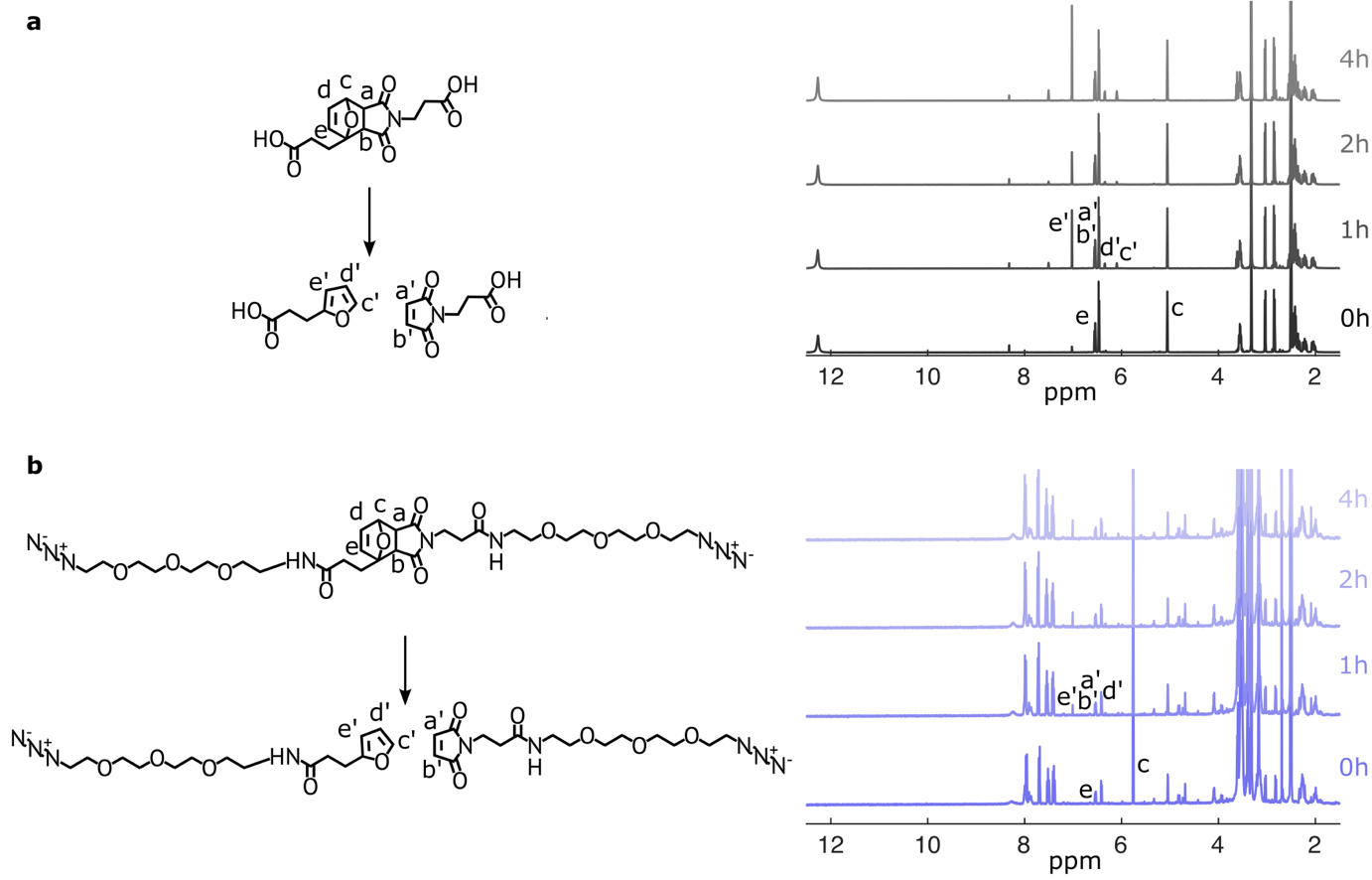

Fig. S9. (a) Conceptional illustration of the rDA reaction of the DA adduct at 70 °C yielding the decomposition products, and the ^1^H NMR spectra monitoring the reaction. (b) Conceptional illustration of the rDA reaction of the ThermoXlinker at 70 °C yielding the decomposition products, and the ^1^H NMR spectra monitoring the reaction.

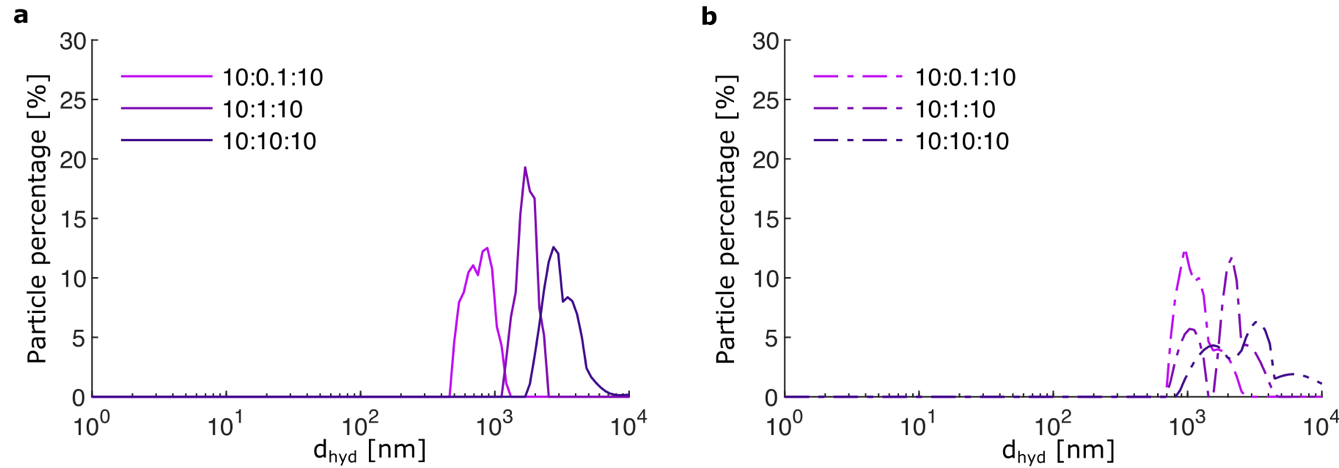

Fig. S10. Hydrodynamic diameters of particle assemblies formed with different ratios between the reactants (particle concentration mg ml^-1^: linker concentration mg ml^-1^: ml solution). (a) Alkyne-based assemblies (n=3). (b) DBCO-based assemblies (n=3).

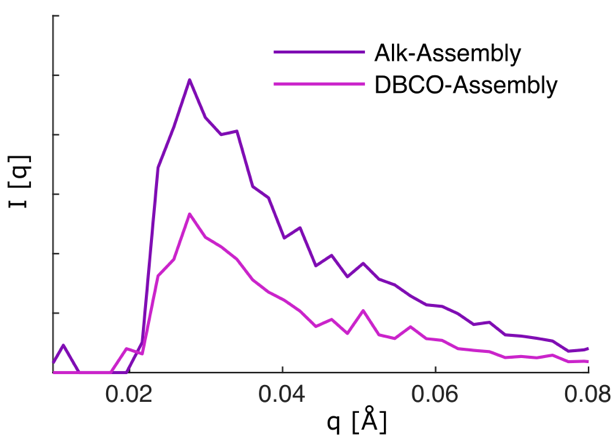

Fig. S11. SAXS spectra of alkyne(Alk)-based assemblies and DBCO-based assemblies

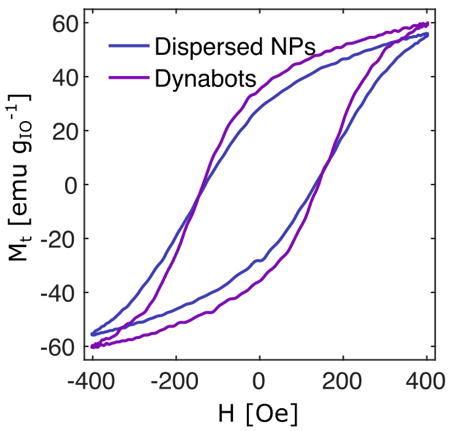

Fig S12: Simulated AC hysteresis loops and SAR for single zinc substituted iron oxide particles and assembled Dynabots. The dynamic magnetic properties are almost identical. The inset shows the assumed distribution of MNPs within the assembly used for simulating the AC magnetic loops of Dynabots, with an interparticle distance of 41 nm (15 + 26/2 nm).

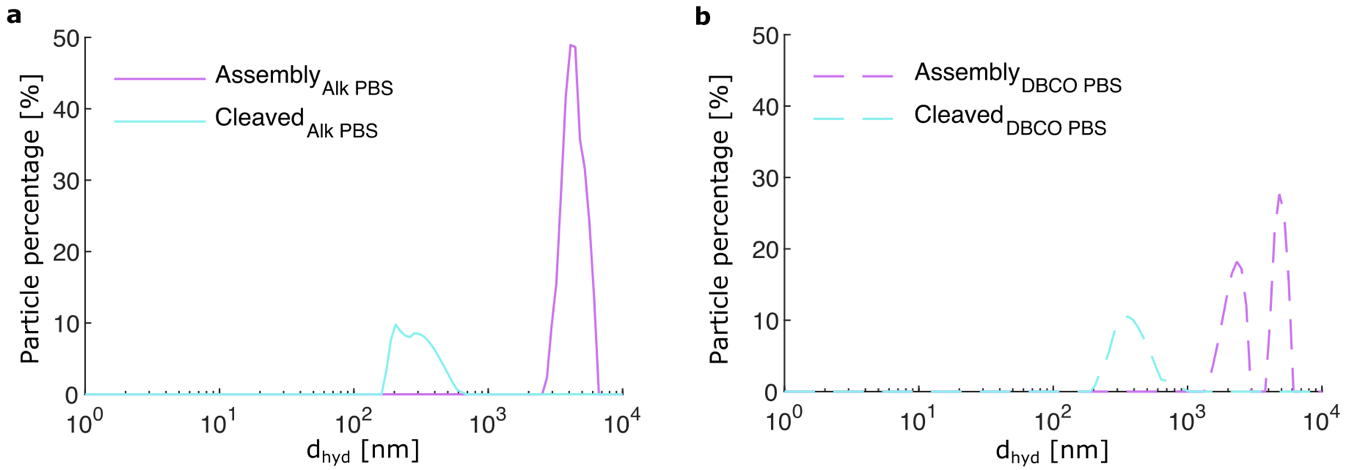

Fig. S13. DLS spectra of assemblies and building blocks of: (a) Alk-based assemblies in PBS after 3 weeks (n=3). (b) DBCO-based assemblies in PBS after 3 weeks (n=3).

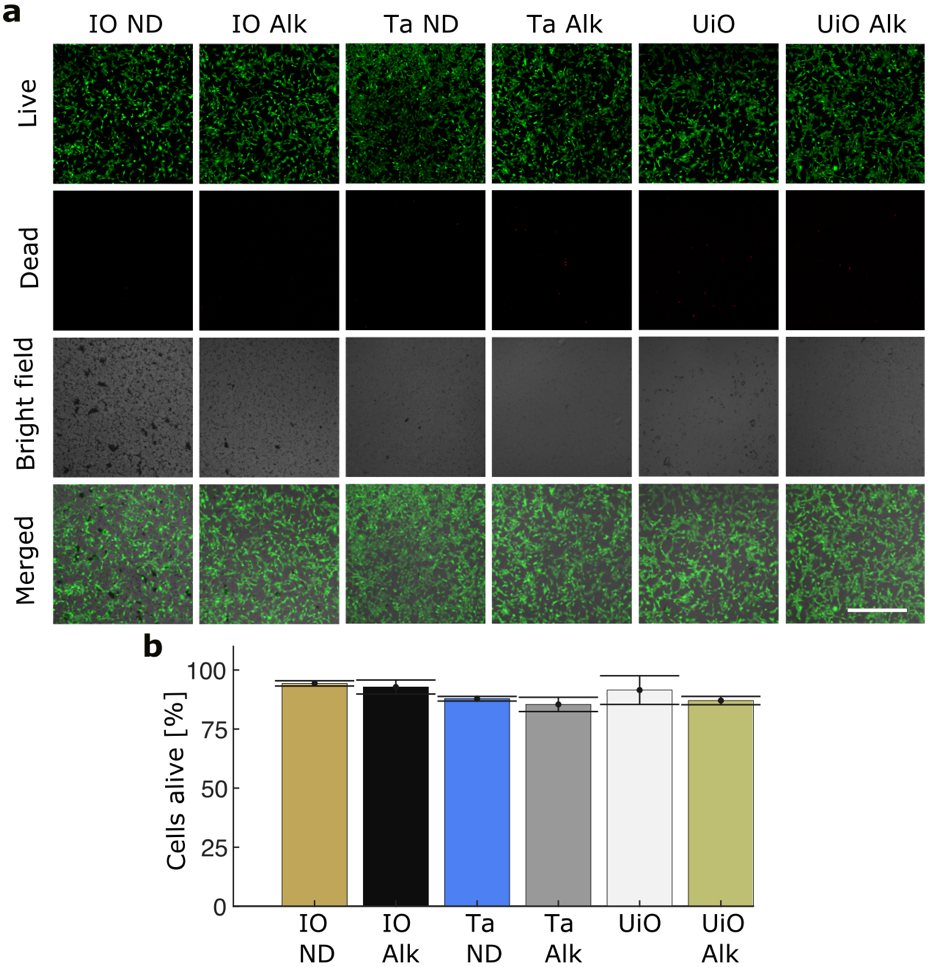

Fig. S14. (a)-(b) Live-dead assays of cells exposed to nitrodopamine-functionalized zinc-substituted iron oxide nanoparticles (brown); alkyne functionalized zinc-substituted iron oxide nanoparticles (black); nitrodopamine functionalized tantalum nanoparticles (brown); alkyne functionalized zinc-substituted iron oxide nanoparticles (black); UiO_66_-NH_2_ (white); UiO_66_- NH_2_ alkyne (yellow).

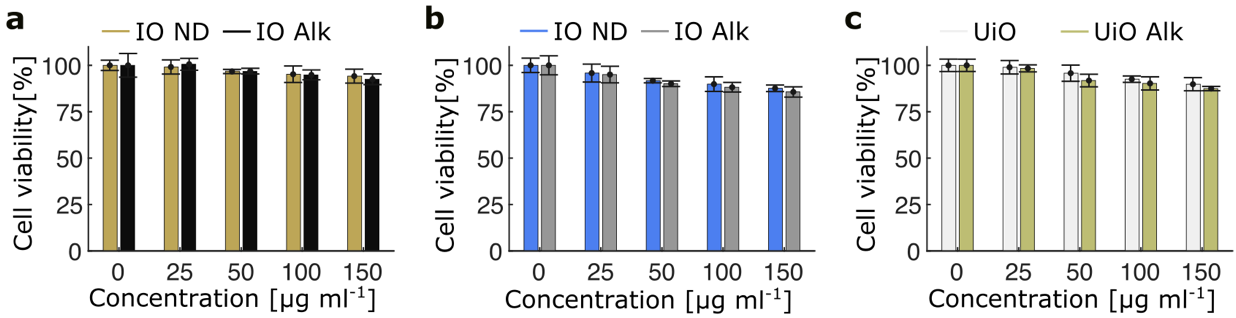

Fig. S15. MTT assays performed on NIH/3T3 cells exposed to: (a) zinc-substituted iron oxide nanoparticles. (b) tantalum nanoparticles. (c) UiO-66-NH_2_ nanoparticles

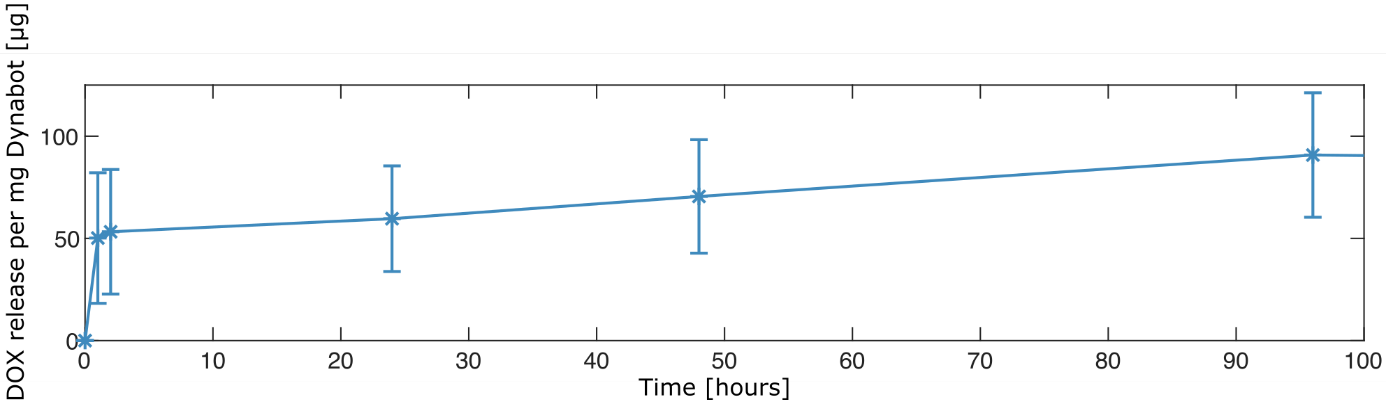

Fig. S16. Doxorubicin release profile from Dynabots at a pH 5.5 in PBS.

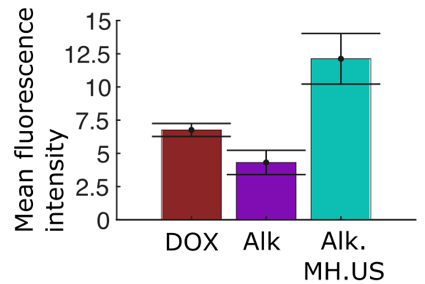

Fig. S17. Comparison of fluorescence intensity of 3D B16-F10 cell spheroid after incubation with DOX, DOX loaded Alk. based assemblies for 4 h with and without magnetic hyperthermia and ultrasound stimulation

To assess whether the core temperature at the Dynabot-rich region could exceed the ThermoXlinker activation threshold, we performed finite-element thermal modelling using COMSOL Multiphysics. The peritumorally injected Dynabot suspension was modeled as a dispersed, disc-shaped heat-generating region embedded beneath the skin, reflecting the expected spreading of the injected suspension. Under the experimental 808 nm irradiation condition, the model predicted a steep spatial thermal gradient around the heat-generating region. The local temperature within the Dynabot-rich region reached 72 °C, whereas the temperature at the tissue surface remained 55–60 °C. These results suggest that infrared thermography underestimates the local temperature experienced by the Dynabot assemblies and supports the feasibility of ThermoXlinker cleavage under the applied in vivo irradiation conditions.

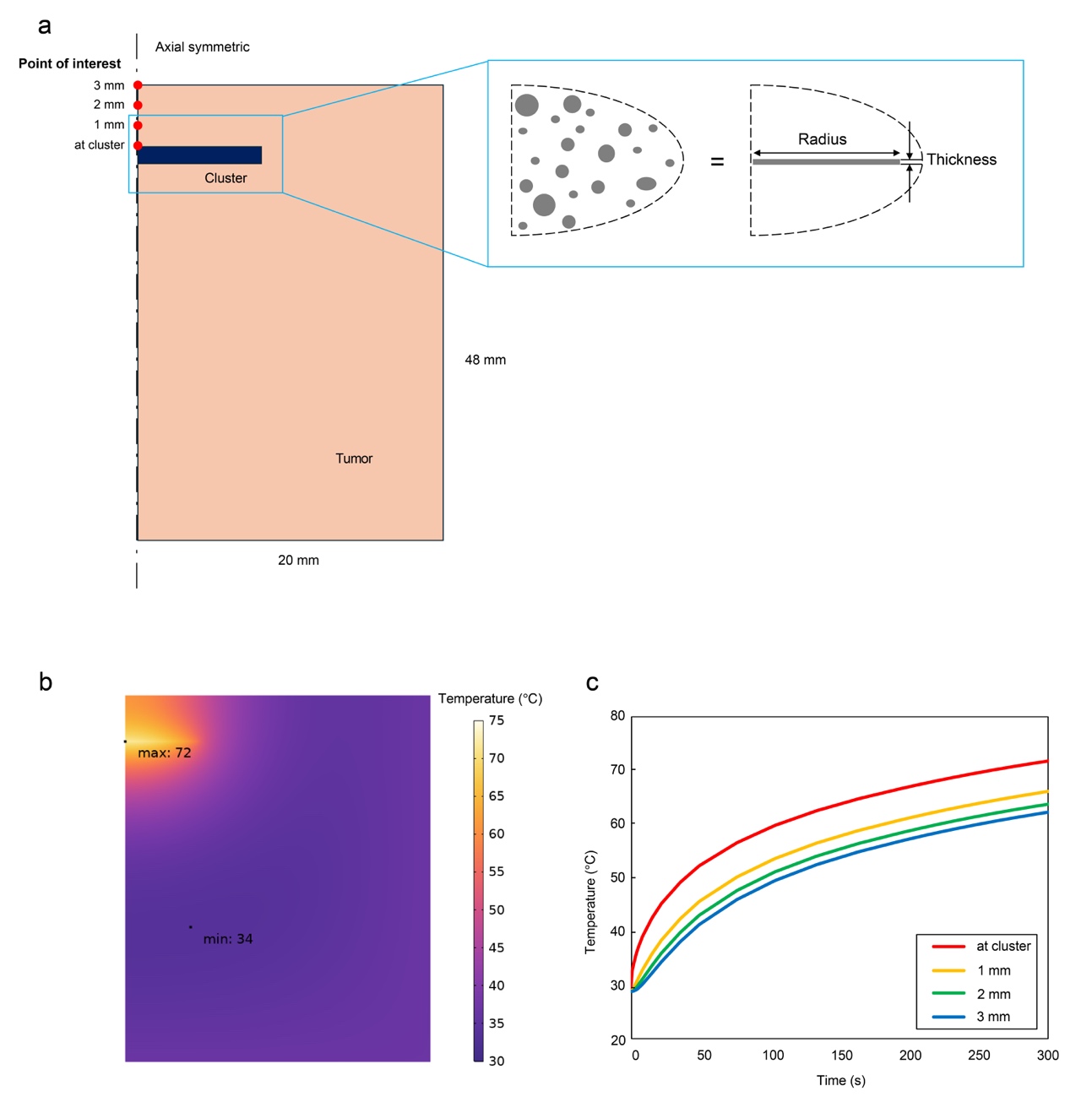

**Fig. S18.** COMSOL-based estimation of local Dynabot temperature under in vivo NIR irradiation. (a) Schematic of the finite-element model, in which the peritumorally injected Dynabot suspension was represented as a dispersed disc-shaped heat-generating region beneath the skin. (b) Representative simulated temperature distribution under 808 nm irradiation. (c) Simulated temperature profiles at different tissue depths

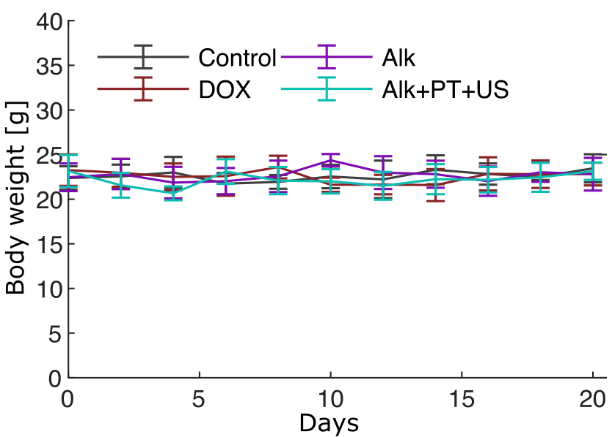

Fig. S19. Body weights of rodent specimens throughout the treatment duration (n=5).

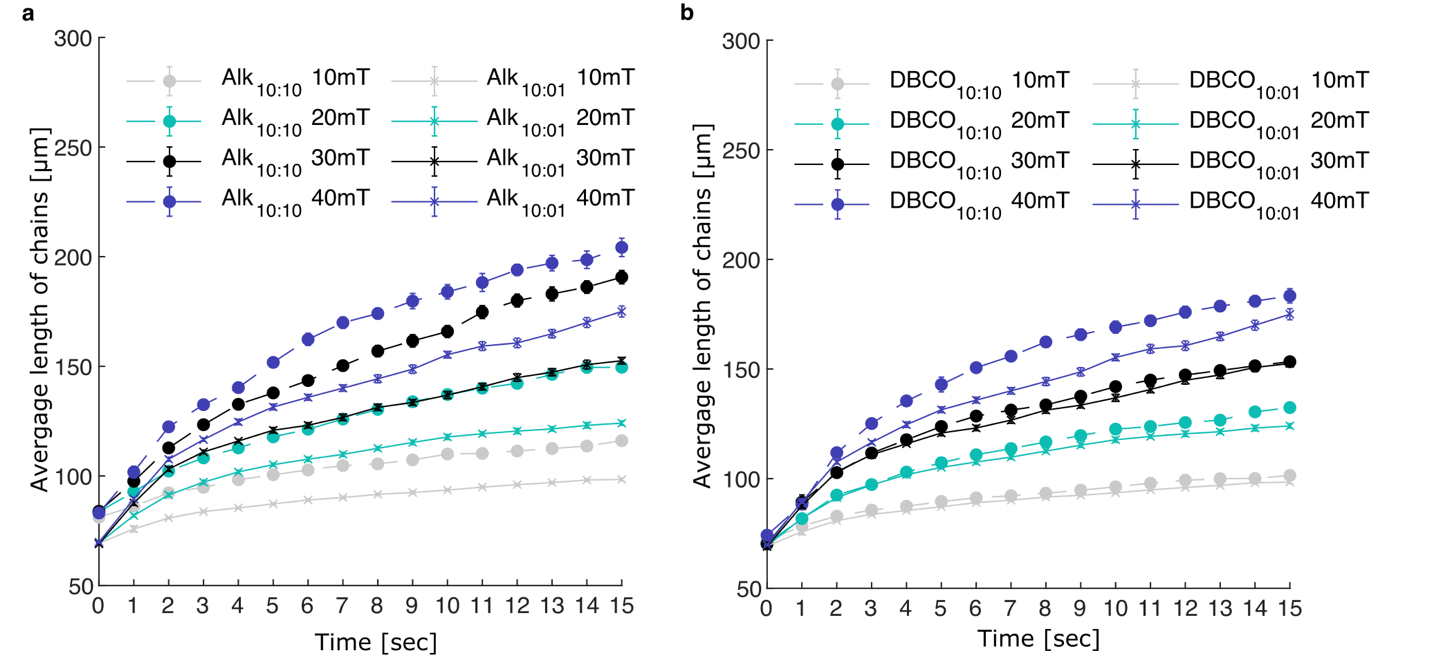

Fig. S20. Swarm arrangement development in dependency of applied field strength. (a) Alkyne based assemblies. (b) DBCO based assemblies.

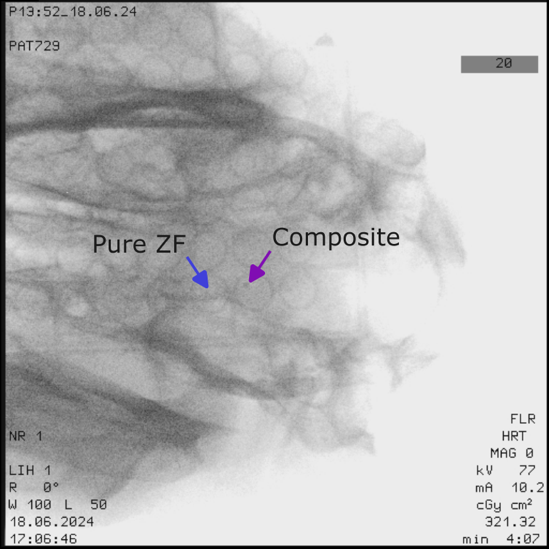

Fig. S21. Ex vivo x-ray fluoroscopy image of monolithic Dynabots comprising pure zinc-substituted ferrite nanoparticles (Pure ZF) and a mixture of ZF, tantalum, and UiO_66_-NH_2_ nanoparticles (Composite) under an ovine head.
